# Riboflavin drives nucleotide biosynthesis and iron-sulfur metabolism to promote acute myeloid leukemia

**DOI:** 10.1101/2025.06.26.661633

**Authors:** Stefan Bjelosevic, Rahel Fauth, Brian T. Do, Gabriela Alexe, Lucy A. Merickel, Thomas D. Jackson, Allen T. Basanthakumar, Audrey Taillon, Silvi Salhotra, Birgitta Ryback, Constanze Schneider, Giulia DiGiovanni, Ryan Elbashir, Nils Burger, Muhammad Bin Munim, Shan Lin, Pere Puigserver, David E. Root, Matthew G. Vander Heiden, Kimberly Stegmaier

## Abstract

Riboflavin is a diet-derived vitamin in higher organisms that serves as a precursor for flavin mononucleotide and flavin adenine dinucleotide, key cofactors that participate in oxidoreductase reactions. Here, using proteomic, metabolomic and functional genomics approaches, we describe a specific riboflavin dependency in acute myeloid leukemia and demonstrate that, in addition to energy production via oxidative phosphorylation, a key biological role of riboflavin is to enable nucleotide biosynthesis and iron-sulfur cluster metabolism. Genetic perturbation of riboflavin metabolism pathways or exogenous depletion in physiological culture medium induce nucleotide imbalance and DNA damage responses, as well as impair the stability and activity of proteins which utilize [4Fe-4S] iron-sulfur clusters as cofactors. We identify a window of therapeutic opportunity upon riboflavin starvation or chemical riboflavin metabolism perturbation and demonstrate that this strongly synergizes with BCL-2 inhibition. Our work identifies riboflavin as a critical metabolic dependency in leukemia, with functions beyond energy production.

## Introduction

Cofactors, many of which are derived from vitamins, are organic molecules that facilitate chemical reactions by binding to inactive apoenzymes to enable their function (holoenzymes) and thus drive diverse biological processes^1^. These processes include transfers of functional groups (e.g., acyl, one-carbon and amino groups), electron transfer in redox reactions^2,3^, and enzymes that catalyze epigenetic processes^4^. As such, vitamins and their cognate cofactors play critical roles in cell proliferation and differentiation. While they have been extensively studied in non-malignant contexts, how cofactors and vitamins influence oncogenic transformation and maintenance is surprisingly understudied.

Humans and other eukaryotes have long lost the ability to *de novo* synthesize vitamins and thus they must be acquired through the diet^5^. As a result, evolutionarily acquired mechanisms to store or salvage vitamins or their coenzyme products in periods of dietary restriction have developed. For example, fat-soluble vitamins (A, D, E and K) are readily stored in the liver and other fatty tissues, a phenomenon that may result in toxic accumulation if intake is beyond physiological requirements^6^. Deprivation of the water-soluble B-complex and C vitamins, while less readily stored *in vivo*, can be tolerated for several weeks to months. The onset of clinical manifestations of B-vitamin deficiency (such as thiamine,riboflavin and folate) can take as long as four to five weeks^7–9^, while deficiency of vitamin B_12_ (cobalamin) can be tolerated for several years^10^. Additionally, the evolution of intracellular salvage pathways (for example, the recycling of niacinamide (vitamin B_3_)-derived nicotinamide (NAM) into nicotinamide adenine dinucleotide (NAD+) through nicotinamide phosphoribosyltransferase (NAMPT) catalysis^11^) can ensure the availability of vital coenzymes for enzymatic function during periods of dietary deficiency. These observations raise the possibility that, in the context of cancer, normal tissues can maintain vitamin homeostasis by drawing on physiological reserves, which may be insufficient for the heightened metabolic demand of cancer cells. Thus, dietary deprivation of vitamins could be employed as a therapeutic modality (potentially in combination with other antineoplastic agents), while limiting effects on healthy tissues.

Indeed, the therapeutic targeting of vitamin processing pathways in hematological malignancies, such as acute myeloid leukemia (AML) and acute lymphoblastic leukemia (ALL), has long been employed. The most prolific is the use of folate antimetabolites such as methotrexate and pemetrexed as essential components of chemotherapy regimens used to treat ALL^12,13^. Similarly, acute promyelocytic leukemia (APL), a subtype of AML typically characterized by recurrent *PML::RARA* fusions, is effectively treated by the use of all-*trans* retinoic acid, a vitamin A derivative, in combination with arsenic trioxide^14,15^. Vitamin involvement in leukemogenesis has been previously described; for instance, ascorbate (vitamin C) depletion can impair the function of the ten-eleven translocation (TET) methylcytosine dioxygenase TET2 and cooperate with FMS-like tyrosine kinase 3 (FLT3) mutations to drive AML^16–18^. Targeting of pyridoxine (vitamin B_6_) and its intracellular processing pathways has also been demonstrated to be an AML-specific dependency, as identified by genomic screens^19^. These clinical and experimental observations provide the conceptual impetus for further systematic identification of vitamins and their processing pathways for anti-leukemic intervention.

Herein, we describe an effort to systematically identify vitamin dependencies in AML that can be leveraged for therapeutic gain. Using Plasmax, a liquid culture medium designed to physiologically mimic human plasma^20^, we performed vitamin depletion screens in genetically diverse AML cell lines. We identify vitamin B_2_ – riboflavin – as a potent leukemic dependency. Leveraging the Broad Institute’s Cancer Dependency Map (DepMap), in addition to our own *in vitro* and *in vivo* CRISPR-Cas9 screens, we show that dietary deprivation of riboflavin, or genetic and chemical targeting of its processing pathways, results in collapse of oxidative phosphorylation due to the structural disassembly of mitochondrial complex I and II. We demonstrate that riboflavin is specifically necessary for pyrimidine biosynthesis in AML, and perturbation of riboflavin metabolism induces a DNA damage response due to nucleotide imbalance that can be rescued by either attenuating *de novo* purine metabolism or supplementing liquid culture with pyrimidine nucleosides. Unexpectedly, we find that riboflavin loss also impairs the stability and function of proteins which contain iron-sulfur (Fe-S) clusters, and enzymes necessary for riboflavin metabolism are genetic co-dependencies specifically with proteins which bind the [4Fe-4S] Fe-S cluster. Importantly, the dependency on riboflavin and subsequently impaired oxidative phosphorylation upon perturbation of riboflavin metabolism can be exploited by sensitizing AMLs to the BCL-2 inhibitor venetoclax. Collectively, this work describes a mechanistic framework of riboflavin dependency in AML and defines a previously unappreciated interplay between riboflavin, pyrimidine biosynthesis and Fe-S metabolism.

## Results

### Vitamin depletion screens in physiological culture medium identify riboflavin as an AML proliferative dependency

Traditional culture medium is an imperfect recapitulation of physiological conditions, including supraphysiological levels of metabolites that enable rapid proliferation such as glucose and glutamine^21^. Among other metabolites that have poor physiological fidelity are vitamins; some, such as myoinositol, are present in abundances up to 18 times that of physiological levels, while others, such as vitamin C, are completely absent from basic formulations^22^. To address this limitation, we employed Plasmax, a liquid culture medium designed to recapitulate human plasma^20^. Importantly, Plasmax contains vitamins at empirically determined physiological levels in contrast to other physiological media^23^. In order to evaluate the essentiality of vitamins in the maintenance of the leukemic state, we used Plasmax as a platform to perform vitamin depletion screens in AML cells. We selected two genetically and biologically distinct AML cell lines to perform these experiments: NB4, an APL cell line which contains a *PML::RARA* fusion and MOLM-13, a myelomonocytic cell line which harbors the *KMT2A::MLLT3* (more commonly known as the *MLL-AF9*) fusion, and a FLT3 internal-tandem duplication (*FLT3-ITD*). Both cell lines proliferated continuously and in a comparable manner in Plasmax and RPMI-1640 supplemented with 10% fetal bovine serum until day 6 when proliferation sharply diverged (Figure S1A). We thus ensured that media changes were employed to alleviate nutrient exhaustion in the subsequent screen. To perform the screen, we cultured NB4 and MOLM-13 cells in Plasmax complete medium or Plasmax complete medium supplemented with all-*trans* retinoic acid (ATRA), a clinically employed anti-leukemia vitamin derivative which acted as the positive control. We then systematically depleted vitamins in Plasmax medium for six days and utilized flow cytometry to assess proliferation changes using CountBright Absolute Counting beads or induction of myeloid differentiation using CD11b cell surface expression (Figure 1A, see Methods for media formulations). Medium was refreshed at day 3, where no biologically significant proliferation changes were observed in vitamin depleted cultures (Figure S1B). At day 6, both cell lines showed strong growth inhibition and induction of differentiation in ATRA-treated control cells; only depletion of the vitamins folate and riboflavin induced proliferation loss in both cell lines (Figure 1B). As previously reported, depletion of myo-inositol impaired proliferation in MOLM-13^24^. While folate depletion induced differentiation in both cell lines as measured by increased CD11b surface expression (Figures 1C and 1D), deprivation of riboflavin had negligible effects on CD11b levels. Folate and its downstream processing pathways have long been targeted in leukemia. We thus concentrated further efforts on characterizing the role of riboflavin in AML as this has not been previously explored to our knowledge. We confirmed that riboflavin deprivation induced strong Annexin V-positive cell death in both cell lines used in our original screen and in additional AML cell lines, MV4-11 and OCI-AML2 (Figure 1E). These vitamin depletion screens therefore identify riboflavin as a potent dependency required for AML proliferation in physiological conditions.

**Figure 1.**
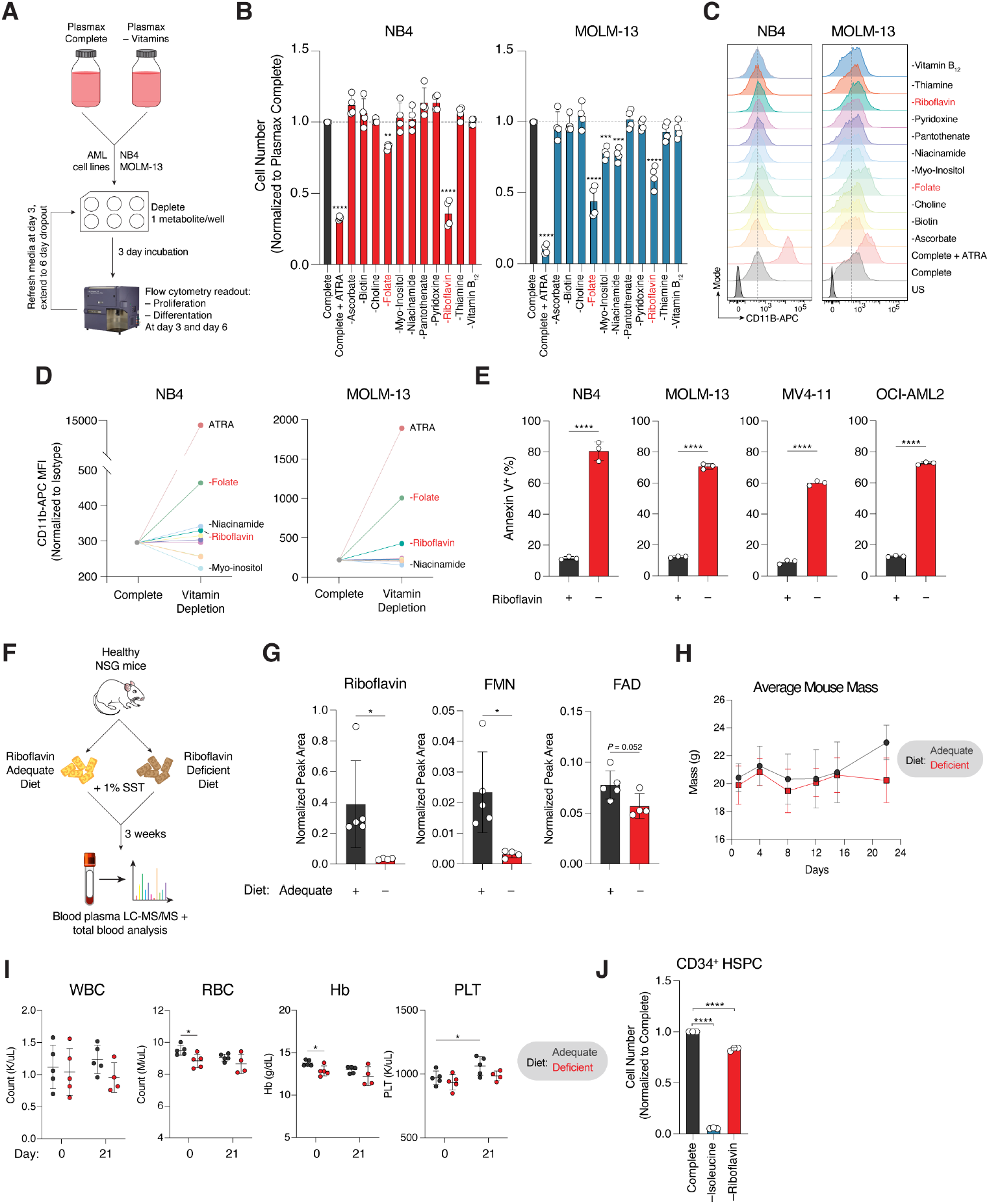
Physiologically relevant metabolite depletion screens identify riboflavin as a leukemia dependency. **A**. Schema of vitamin depletion screens in AML cells. **B**. Proliferation of NB4 and MOLM-13 cells upon systematic removal of vitamins from Plasmax medium at day 6. Cell number is normalized to Plasmax complete medium. Cells cultured in Plasmax complete medium and treated with 1μM all-*trans* retinoic acid (ATRA) served as an anti-proliferation/differentiation control. Data representative of *n*=4 pooled replicates from two independent biological experiments. **C**. Representative histograms of cell surface CD11b-APC in NB4 and MOLM-13 cells upon systematic removal of vitamins from Plasmax medium at day 6. **D**. Quantification of CD11b geometric mean fluorescence intensity (MFI) in NB4 and MOLM-13 cells from C. at day 6. The average of 2 replicates per condition was normalized to the isotype control and compared to Plasmax complete. **E**. Cell death in NB4, MOLM-13, MV4-11 and OCI-AML2 cells as measured by Annexin V-APC cell surface staining at day 6 (NB4, OCI-AML2), day 8 (MV4-11) and day 11 (MOLM-13) post riboflavin withdrawal from Plasmax. Data representative of *n*=3 biological replicates. **F**. Schema of riboflavin depletion experiments in NSG mice. SST, succinylsulfathiazole; LC-MS/MS, liquid chromatography with tandem mass spectrometry. **G**. Quantification of riboflavin, FMN and FAD in blood plasma of mice fed adequate (*n*=5) versus deficient (*n*=4) riboflavin diets at 21 days. **H**. Average mouse mass of mice fed adequate (*n*=5) versus deficient (*n*=4) riboflavin diets over 21 days. **I**. Complete blood counts of mice fed adequate versus deficient riboflavin diets at start of experiment (baseline, day 0) and at 21 days. WBC, white blood cell count; RBC, red blood cell count; Hb, hemoglobin test; PLT, platelet count. **J**. Cell number of healthy CD34^+^ HSPC cells after 6 days of culture in Plasmax complete or Plasmax lacking isoleucine (anti-proliferative control) or riboflavin. Cell number is normalized to Plasmax complete medium. Data representative of *n*=3 biological replicates. The experiment was repeated with two separate donors. Data are presented as the mean ± SD. \**P* < 0.05; \*\**P* < 0.01; \*\*\**P* < 0.001 and \*\*\*\**P* < 0.0001 by ordinary one-way ANOVA with Bonferroni’s multiple comparisons test (B, J) and unpaired two-tailed Student’s t-test (E, G, I).

To test whether normal hematopoiesis can tolerate periods of dietary riboflavin starvation *in vivo*, we placed non-tumor bearing NOD-*scid* IL2Rg^null^ (NSG) mice on a riboflavin competent (0.006 g/kg) or deficient diet for 3 weeks (Figure 1F). Given that gut microflora retains the ability to *de novo* synthesize riboflavin^25^, we attempted to control for this phenomenon by including the antibiotic succinylsulfathiazole (SST) in both diets. We reasoned that this inclusion retained clinical relevance, as prophylactic antibiotics are employed in frontline AML therapy regimens^26^. We isolated blood plasma from mice at endpoint and performed liquid chromatography with tandem mass spectrometry (LC-MS/MS)-based metabolite profiling for riboflavin and its downstream metabolites. As expected, plasma riboflavin levels were strongly reduced in mice fed a riboflavin deficient diet, as were levels of the riboflavin cofactor products, flavin mono-nucleotide (FMN) and flavin adenine dinucleotide (FAD) (Figure 1G). These results are consistent with long-appreciated re-distribution of plasma and intracellular flavin pools to preserve FAD at the expense of FMN and riboflavin, likely an ancient evolutionarily evolved mechanism to maintain critical cellular functions, such as respiration, in periods of dietary starvation^27,28^. Mice starved of dietary riboflavin did not significantly lose body mass (Figure 1H). Whole-blood analysis of mouse hematopoiesis did not demonstrate any significant reductions in white blood cells (WBC), red blood cells (RBC), hemoglobin (Hb) or platelets (PLT), among other parameters (Figure 1I and S1C), in riboflavin deficient mice. To further test the hypothesis that AML cells have heightened requirements for riboflavin, we cultured healthy donor-derived human CD34^+^ hematopoietic stem and progenitor (HSPC) cells in Plasmax complete medium, in Plasmax lacking riboflavin, or in Plasmax lacking the essential amino acid isoleucine as a positive control. At day 6, riboflavin loss had mild impacts on HSPC proliferation, while in AML cells, riboflavin loss induced cell death. Isoleucine deprivation completely ablated proliferation in these cells (Figure 1J). While short term culture in riboflavin free medium did not impact normal HSPCs, extended culture longer than 14 days reduced HSPC proliferation by approximately 60% (Figure S1D). Thus, these experiments support the potential for an exploitable therapeutic window for riboflavin starvation in AML cells versus untransformed hematopoietic cells.

### Riboflavin kinase is a metabolic vulnerability in AML

Dietary-derived riboflavin is primarily absorbed in the proximal small intestine or in the blood via three dedicated membrane transporters, SLC52A1, SLC52A2 and SLC52A3^29–31^. Upon intracellular uptake, riboflavin is phosphorylated by riboflavin kinase (RFK) to yield FMN, a cofactor which participates in oxidoreductase, dehydrogenase and monooxygenase reactions^32^. FMN is subsequently adenylated by flavin adenine dinucleotide synthetase 1 (FLAD1) to form FAD, a cofactor involved in similar processes (Figure 2A). Importantly, FMN and FAD are hydrophilic and unable to cross the plasma membrane; they must first be sequentially hydrolyzed by the non-specific nucleotidase CD73 and alkaline phosphatase to riboflavin to facilitate entry^33^. To investigate the importance of these riboflavin processing enzymes in cancer, we mined the Broad Institute’s Cancer Dependency Map (DepMap), a collection of over 1,100 genome-wide CRISPR-Cas negative selection screens across a wide range of human cancers. Strikingly, *RFK*, the first and rate-limiting enzyme of riboflavin metabolism, was a strongly enriched dependency in hematological malignancies (myeloid and lymphoid) (Figure 2B). While mRNA expression of *RFK* in myeloid and lymphoid cell lines was significantly less than most solid cell lines (Figure S2A), *RFK* dependency did not correlate with mRNA expression (Figure S2B). To functionally validate the dependency on *RFK* in AML, we performed *in vitro* competition experiments in five genetically diverse AML cell lines by transducing a doxycycline-inducible CRISPR single-guide RNA (sgRNA)-GFP construct targeting *RFK* into cell lines bearing Cas9-mCherry. As expected, *RFK* loss potently suppressed AML proliferation and led to a strong loss of representation in *RFK* knockout (KO) cells as measured by decreased GFP positivity and compared to a control sgRNA targeting the conserved *ROSA26* locus (sgRosa) (Figures 2C, 2D and S2C). We further confirmed *RFK* dependency in two Cas9-competent orthotopic patient-derived xenograft (PDX) models, PDX16-01 and PDX17-14, which are capable of short-term growth in liquid culture^24^ (Figure 2E and S2D). A vast majority of hematological cancer cell lines screened in DepMap are cultured in RPMI, and absolute riboflavin concentrations are comparable between RPMI and Plasmax (500 nM versus 300 nM). Nevertheless, to rule out the possibility that micronutrients exclusively present in plasma/physiological medium can compensate for or circumvent perturbation of riboflavin metabolism, we confirmed that *RFK* deletion in AML cells cultured in Plasmax phenocopied the effects of culture in RPMI, suggesting that this dependency is relevant in physiological systems (Figure 2F). Importantly, ectopic overexpression of a wild-type (WT) *RFK* cDNA engineered to be sgRNA resistant completely rescued the antiproliferative effect of RFK loss, including in PDX16-01 cells cultured *in vitro* (Figure 2G). In agreement with our dependency analysis suggesting solid cancer cell lines are largely unaffected by *RFK* loss, deleting *RFK* or exogenously depleting riboflavin in Plasmax culture in two solid cancer models, EW8 (Ewing sarcoma) and SKNAS (neuroblastoma), had no effect on cell proliferation (Figure 2H, S2E and S2F).

**Figure 2.**
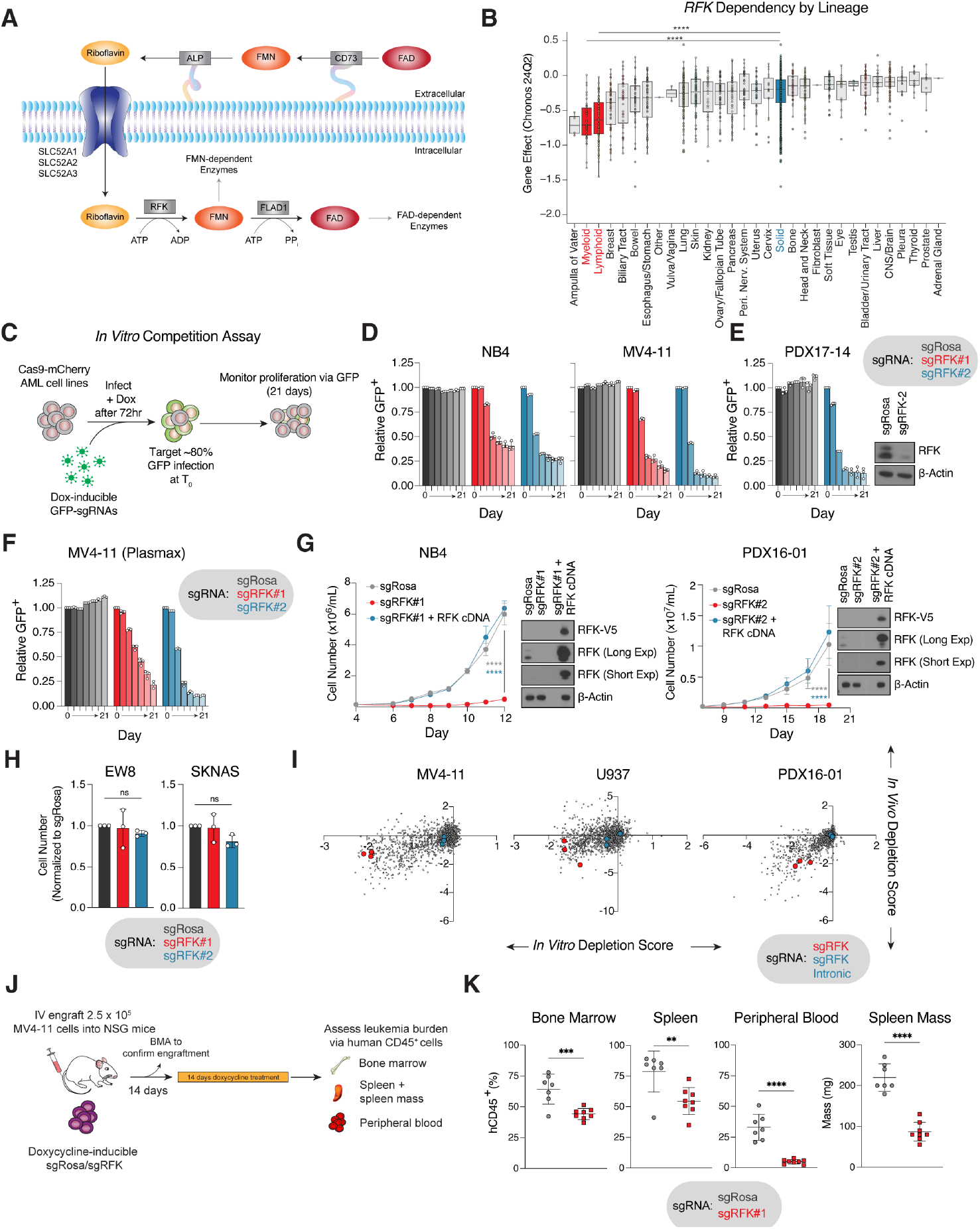
Riboflavin kinase (RFK) is a metabolic dependency in acute myeloid leukemia. **A**. Schema of cellular riboflavin processing pathways. FMN, flavin mononucleotide; FAD, flavin adenine dinucleotide; ALP, alkaline phosphatase; RFK, riboflavin kinase; FLAD1, flavin adenine dinucleotide synthase 1. **B**. Dependency on RFK in cancer cell lines by disease lineage from the Broad Institute’s DepMap 24Q2 database. Gene essentiality reported as Gene Effect Chronos score, where more negative values equal higher essentiality and each data point represents an individual cell line. Myeloid and lymphoid lineage are highlighted in red, pooled solid cancer cell lines in blue. Boxplots ranked by median RFK dependency. **C**. Schema of in vitro competitive proliferation assays. **D**. Competitive proliferation experiments in NB4 and MV4-11 AML cell lines over 21 days in RPMI medium. Two independent *RFK* sgRNAs and a control targeting the conserved *ROSA26* locus (sgRosa) were used, and relative proliferation was measured using GFP as a readout of edited cells. Data representative of *n*=3 biological replicates. **E**. Competitive proliferation experiments in Cas9-competent PDX17-14 cells subjected to short-term *in vitro* culture over 21 days in PDX medium. An *RFK* sgRNA and a control targeting Rosa was used, and relative proliferation was measured using GFP as a readout of edited cells. Data representative of *n*=3 biological replicates. Western immunoblot of RFK KO shown, with β-actin serving as the loading control. **F**. Competitive proliferation experiments in the MV4-11 AML cell line over 21 days in Plasmax complete medium. Two independent RFK sgRNAs and a control (sgRosa) were used, and relative proliferation was measured using GFP as a readout of edited cells. Plasmax medium was refreshed every 72 hours. Data representative of *n*=3 biological replicates. **G**. Proliferation of NB4 or PDX16-01 cells upon induction of a doxycycline-inducible sgRNA targeting sgRosa, RFK, or endogenous RFK knockout cells expressing a wild type (WT) CRISPR-resistant *RFK* cDNA construct over 12 days (NB4) or 19 days (PDX16-01). Data representative of *n*=3 biological replicates. Validation western immunoblots of endogenous RFK and V5-tagged RFK over-expression and knockout shown, with β-actin serving as the loading control. **H**. Cell number of EW8 Ewing sarcoma and SKNAS neuroblastoma cells upon genetic deletion of *RFK* using two independent sgRNAs and a control (sgRosa) at 14 days. Cell number is normalized to sgRosa. Data representative of *n*=3 biological replicates. **I**. Scatter plots of the *in vitro* and *in vivo* depletion scores of RFK at the gene level in MV4-11, U937 and PDX16-01. Data points representing the median value of *RFK* (red circles) and its matching intronic control (blue circles) are shown. **J**. Schema of *in vivo* experiment examining the anti-leukemic effects of *RFK* loss in MV4-11 cells. **K**. Examination of key hematological compartments at 14 days post RFK depletion *in vivo*. Data reported as percentage of human CD45-positive cells of total cells. Data are presented as the mean ± SD. \*\*\**P* < 0.001 and \*\*\*\**P* < 0.0001 by ordinary one-way ANOVA with Bonferroni’s multiple comparisons test (B, H), ordinary two-way ANOVA with Bonferroni’s multiple comparisons test (G), and individual unpaired two-tailed Student’s t-test (K).

To determine whether targeting riboflavin processing has *in vivo* antileukemic efficacy, we took advantage of a previous *in vivo* CRISPR screening platform established in our laboratory, which employed a 200 gene library enriched for AML dependencies, and which included sgRNAs targeting *RFK*^24^. In all AML models tested, including PDX16-01, sgRNAs targeting *RFK* were strongly depleted both *in vitro* and *in vivo*, while control sgRNAs targeting *RFK* intronic sequences were not (Figure 2I). We orthogonally validated these results by engrafting Cas9-competent MV4-11 AML cells transduced with a Tet-inducible, GFP-positive sgRNA targeting *RFK* into NSG mice. After confirming engraftment via bone marrow aspiration at day 14, we induced guide activity via introduction of doxycycline-containing chow (625 ppm) for 14 days (Figure 2J). Analysis of bone marrow, spleen and peripheral blood showed marked reduction of human CD45-positive blasts and a concomitant reduction in splenomegaly (Figure 2K), confirming that *RFK* is an AML dependency *in vivo*.

To determine the functional consequence of *RFK* loss, we performed apoptotic and differentiation assessments in our AML models. *RFK* KO induced cell death as measured by Annexin V-positivity (Figure S2G). Interestingly, genetic *RFK* ablation induced differentiation in multiple AML cell lines, including PDX17-14, by flow cytometric CD11b analysis and morphological/histological assessment (Figure S2H and S2I). Collectively, these studies identify the enzyme catalyzing the first step of riboflavin processing, *RFK*, as an AML dependency *in vitro* and *in vivo*, and loss of *RFK* induces potent antiproliferative effects in AML blasts accompanied by differentiation and apoptosis.

### Global proteomics upon perturbation of riboflavin metabolism

To identify proteins regulated by the availability of riboflavin and its cofactors, we performed Tandem Mass Tag (TMT) proteomic analyses in NB4 cells that had undergone *RFK* deletion for 9 days or exogenous riboflavin starvation for 4 days (Figure 3A). We selected these timepoints to enrich for viable cells, as most AML cell lines tested underwent apoptosis by day 11 of genetic *RFK* ablation and day 6 of exogenous riboflavin depletion (Figure S3A). Statistically significant protein abundance changes in *RFK* deletion and exogenous riboflavin starvation were limited (Figure 3B; Table S1) but highly correlated (*R* = 0.511; *P* < 2.2 × 10^−16^) (Figure 3C), with no downregulated proteins unique to exogenous riboflavin depletion but 50 proteins unique to genetic depletion (Figure S3B). This data supports the contention that targeting riboflavin metabolism in AML perturbs a selective and limited number of downstream protein targets.

**Figure 3.**
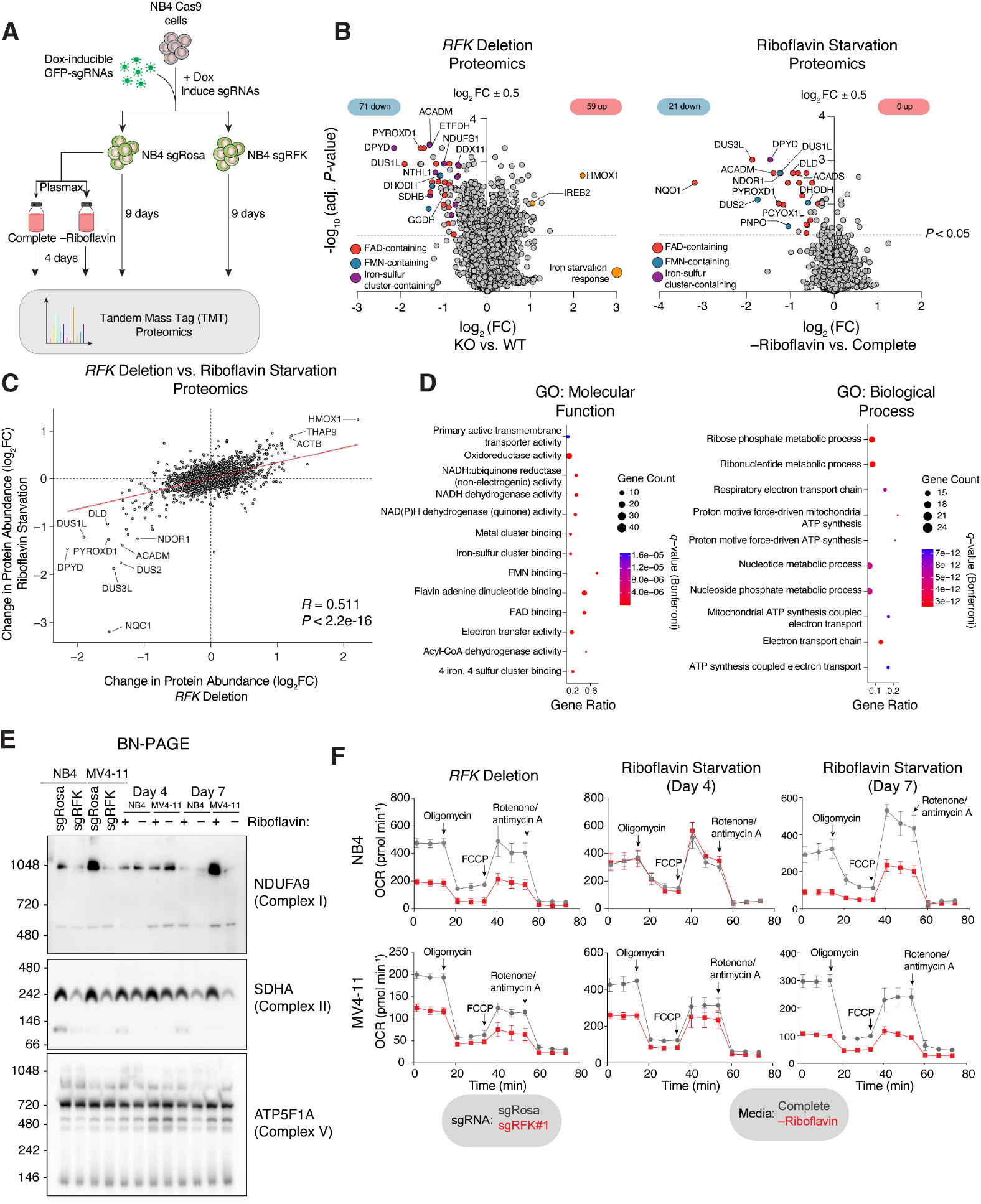
Global analysis of total proteome under riboflavin metabolism perturbation. **A**. Schema of proteomic experiments performed upon genetic depletion of *RFK* (at day 9) or exogenous depletion of riboflavin (at day 4) in NB4 cells. **B**. Volcano plot of relative fold change (log_2_) in protein abundance in NB4 cells upon RFK deletion versus Rosa control at day 9 ranked by significance (left) or riboflavin starvation versus complete medium at day 4 ranked by significance (right). Multiple comparisons were controlled by applying a Benjamini-Hochberg correction (false discovery rate, FDR). The dotted line denotes *P* < 0.05. Red circles, FAD-containing proteins; blue circles, FMN-containing proteins; purple circles, iron-sulfur cluster-containing only proteins; orange circles, iron starvation response-associated proteins. **C**. Pearson correlation plot of the log_2_ change in protein abundance in exogenous riboflavin medium depletion at day 4 versus genetic depletion of *RFK* at day 9 in NB4 cells. Pearson correlation coefficient *R* = 0.511, *P* < 2.2 × 10^−16^. **D**. Bubble plots of enriched gene ontology molecular function (left) and biological process (right) associated with differentially downregulated proteins from *RFK* deletion (B, left pane). *q*-values with Bonferroni correction are reported. **E**. Blue Native (BN) polyacrylamide gel electrophoresis (PAGE) of NB4 and MV4-11 cells at 9 days post deletion of RFK and after 4 and 7 days of exogenous riboflavin starvation. **F**. Oxygen consumption rate (OCR) of NB4 and MV4-11 cells after 9 days of RFK deletion compared to Rosa control, or 4- or 7-days post riboflavin starvation compared to complete medium control. FCCP, carbonyl cyanide 4-(trifluoromethoxy) phenylhydrazone.

Given the well-characterized role of the flavin cofactors FMN and FAD in facilitating electron transfer, particularly within the mitochondrial electron transport chain (ETC)^34^, downregulated proteins in *RFK* KO conditions were strongly enriched for multiple ETC signatures including NADH:ubiquinone reductase, electron transfer activity and proton motive force-driven mitochondrial ATP synthesis by gene ontology (GO) analysis, as well as reduced individual abundance of key proteins such as NDUFV1, a core subunit of mitochondrial complex I^35^ (Figure 3B and 3D). Consistent with previous literature, exogenous depletion of riboflavin at day 4 largely did not impair core ETC subunits, but preferentially depleted proteins which catalyze β-oxidation of medium chain fatty acids, such as the mitochondrial acyl-coenzyme A dehydrogenase ACADM and the glutaryl-CoA dehydrogenase GCDH which catalyzes the oxidative decarboxylation of glutaryl-CoA to crotonyl-CoA^27,36^ (Figure S3B). Unexpectedly, signatures associated with nucleotide metabolism were also represented in our GO analysis as was a reduction in proteins requiring a cognate iron-sulfur (Fe-S) cluster cofactor. Several of these, including NTHL1 (a DNA glycosylase that contributes to DNA repair^37^), SDHB (the iron-sulfur subunit of succinate dehydrogenase^38^) and DDX11 (a helicase involved in DNA repair^39^) require a Fe-S cluster and are not reported to bind FMN or FAD. These results were accompanied by upregulation of IREB2/IRP2, a protein which senses iron depletion and binds to 3’-UTRs of iron-responsive elements to stabilize the mRNA transcripts of genes involved in iron import^40^ and HMOX1, an enzyme which catalyzes the degradation of heme to liberate iron^41^. Many of the other upregulated proteins were associated with chemokine/receptor binding and immune signaling signatures (Figure S3C).

To explore the mechanistic basis by which riboflavin perturbation impairs the ETC in AML, we deleted *RFK* in two AML cell lines (NB4 and MV4-11) or starved AMLs of exogenous riboflavin at two time points (4 and 7 days) and performed Blue Native (BN)-PAGE and SDS-PAGE analysis to assess the assembly and abundance of mitochondrial complexes and individual protein subunits. *RFK* ablation significantly and consistently ablated the levels of fully assembled complex II, and, to a lesser extent, complex I, as did longer-term riboflavin starvation, while other mitochondrial complexes and miscellaneous mitochondrial proteins were largely unaffected (Figure 3E, S3D and S3E). Impaired assembly of mature complex I and II was accompanied by a strongly decreased oxygen consumption rate and maximal respiration capacity upon genetic deletion of *RFK* and a progressive reduction upon riboflavin withdrawal that maximally manifested under longer-term riboflavin starvation (Figure 3F). These data support the contention that *RFK* loss not only ablates canonical electron transport activity in the ETC but is also required for the stability and assembly of mature complex I and II^42^.

### Perturbation of riboflavin metabolism induces nucleotide imbalance

We next sought to determine the metabolic consequences of riboflavin metabolism perturbation. We performed untargeted steady-state LC-MS/MS metabolite profiling which revealed a small number of metabolites as differentially altered upon *RFK* deletion (Figure 4A, Table S2) or acute/chronic riboflavin starvation (Figure S4A, Table S2). Importantly, in both contexts, FMN and FAD were strongly reduced as expected; indeed, even 4 days of riboflavin starvation largely depleted riboflavin and FMN and strongly reduced FAD pools, partially explaining the temporal cell viability differences between genetic *RFK* loss and exogenous depletion (Figure S4A and S4B). Strikingly, differentially altered metabolites upon *RFK* loss or riboflavin starvation were centered around two primary biological processes: nucleotide and nucleotide sugar metabolism (in agreement with our proteomic GO analyses), and the Kennedy Pathway, which converts ethanolamine and choline to phosphatidylethanolamine (PE) and phosphatidylcholine (PC) respectively^43^ (Figure S4C). Namely, upon *RFK* loss and riboflavin starvation, we observed reductions in multiple upstream pathway intermediates, coupled with increase in CDP-choline and CDP-ethanolamine, suggesting incomplete biosynthesis of PE and PC (Figure S4D).

**Figure 4.**
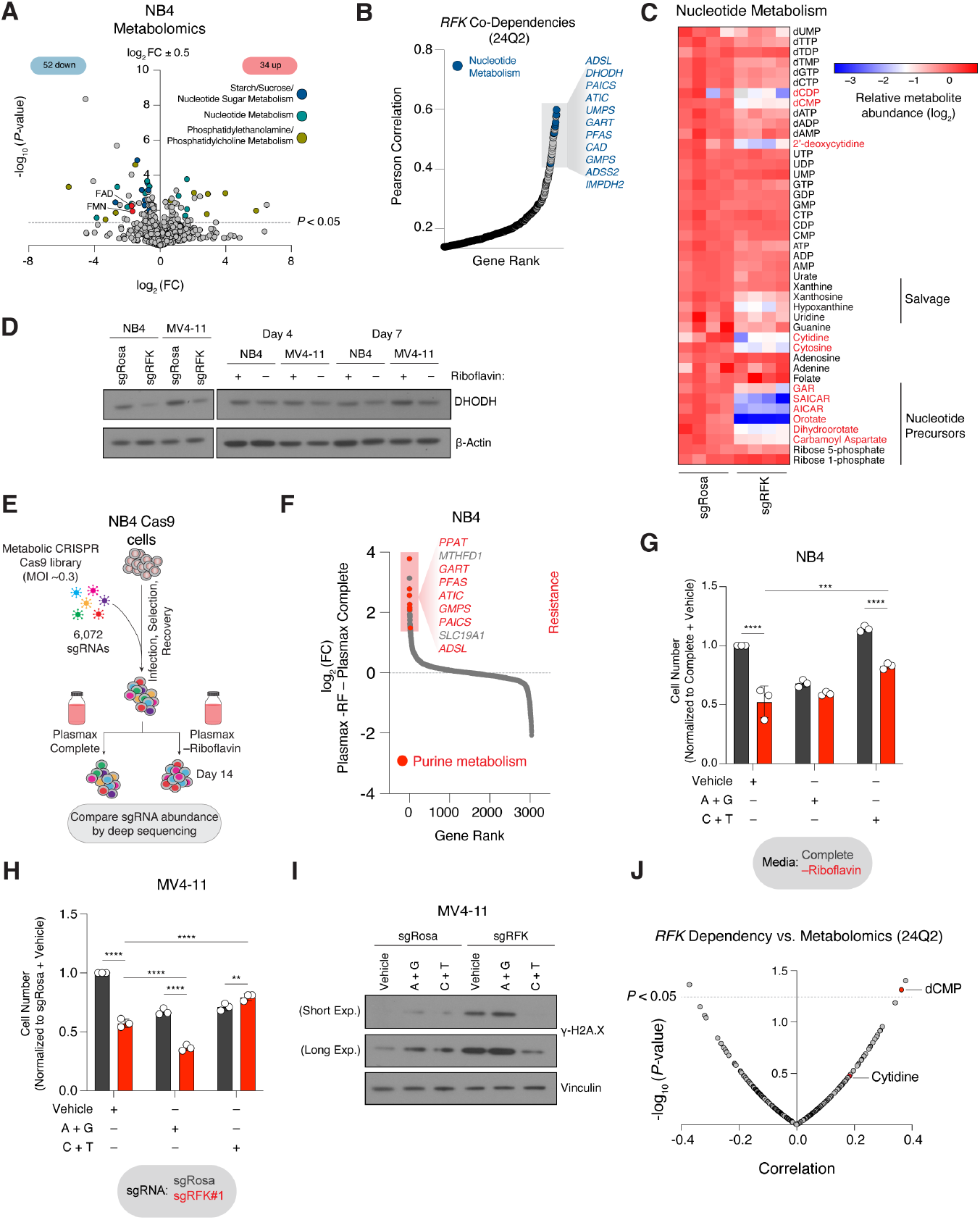
Riboflavin metabolism perturbation induces nucleotide imbalance. **A**. Volcano plot of relative fold change (log_2_) in metabolite abundance in NB4 cells upon RFK deletion versus Rosa control at day 9 ranked by significance. The dotted line denotes *P* < 0.05. Blue circles, starch/sucrose/nucleotide sugar metabolism; green circles, nucleotide metabolism; yellow circles, phosphatidylethanolamine/phosphatidylcholine metabolism. **B**. Dot plot of genetic co-dependency of genes with *RFK* ranked by Pearson correlation. Data from DepMap 24Q2. Blue shading denotes nucleotide metabolism-associated genes. **C**. Heatmap of nucleotide metabolites in *RFK* deleted NB4 cells versus Rosa controls cultured in RPMI for 7 days. Relative log_2_ of metabolite abundances shown. Data representative of *n*=4 biological replicates per condition. **D**. Western immunoblot for DHODH in NB4 and MV4-11 cells upon genetic KO of *RFK* for 9 days (left) or 4 and 7 days of exogenous riboflavin starvation (right). β-Actin served as the loading control. Blot membrane was stripped and re-probed for other proteins of interest, and data from the same gel, including loading control, are also used in Figure S5A. **E**. Schema of CRISPR screen using metabolic library in NB4 cells cultured in Plasmax complete or riboflavin depleted medium. **F**. Dot plot of genes ranked by relative log_2_ enrichment or depletion in Plasmax complete versus riboflavin depleted culture medium as measured by sgRNA abundance. Genes whose abundance increased conferred resistance to riboflavin depletion. Red denotes genes associated with purine biosynthesis. **G**. Cell number of NB4 cells at day 6 post culture in Plasmax complete or lacking riboflavin and treated with vehicle (water), or cocktails of purine (A+G, adenosine + guanosine) or pyrimidine (C+T, cytidine + thymidine) nucleosides. Cell number normalized to Complete + Vehicle. Data representative of *n*=3 biological replicates. **H**. Cell number of MV4-11 cells at day 8 post induction of Rosa or *RFK* sgRNAs, treated with vehicle (water), or cocktails of purine (A+G, adenosine + guanosine) or pyrimidine (C+T, cytidine + thymidine) nucleosides in Plasmax medium for 4 days. Cell number normalized to sgRosa + Vehicle. Data representative of *n*=3 biological replicates. **I**. Western immunoblot for γH2A.X in MV4-11 cells at day 8 post induction of Rosa or RFK sgRNAs, treated with vehicle (water), or cocktails of purine (A+G, adenosine + guanosine) or pyrimidine (C+T, cytidine + thymidine) nucleosides in Plasmax medium for 4 days. Short exposure (top) and long exposure (bottom). Vinculin served as the loading control. **J**. Dot plot of the correlation between *RFK* dependency and metabolite abundance in the DepMap 24Q2 dataset ranked by significance. Each circle denotes an individual metabolite. Dotted line denotes *P* < 0.05. Data are presented as the mean ± SD. \*\**P* < 0.01, \*\*\**P* < 0.001 and \*\*\*\**P* < 0.0001 by ordinary two-way ANOVA with Bonferroni’s multiple comparisons test (G, H).

PE and PC are two key lipid components of the phospholipid bilayer and are involved in other intracellular processes including autophagy and lipid transport^44^. Riboflavin metabolism has not been previously directly linked to the biosynthesis of either metabolite. However, several FAD-dependent acyl-coenzyme A dehydrogenases (ACADs – which are involved in fatty acid oxidation reactions that yield acetyl-CoA, a lipid synthesis precursor) are downregulated upon *RFK* loss and riboflavin starvation. Further orthogonal evidence of the functional role of riboflavin in PE and PC synthesis is provided by the fact that *CHKA*, the first enzyme in the PC pathway which phosphorylates choline to phosphocholine^45^, and *FLVCR1*, the recently identified choline transporter^46^ score as genetic *RFK* co-dependencies, using Cancer Dependency Map (DepMap) data (Figure S4E).

Metabolic outputs of the ETC, such as aspartate and adenosine triphosphate (ATP), are required for nucleotide biosynthesis^47,48^, and FMN/FAD-dependent enzymes are involved in nucleotide biosynthesis processes, such as dihydroorotate dehydrogenase (DHODH), the second enzyme in de novo pyrimidine biosynthesis^49^. Indeed, analysis of *RFK* co-dependencies using DepMap revealed that genes involved in nucleotide metabolism were strongly correlated to *RFK* dependency, consistent with a conserved role for riboflavin in nucleotide biosynthetic processes (Figure 4B). To better understand how riboflavin intersects with nucleotide metabolism, we performed quantitative metabolomics post *RFK* KO and assessed the abundance of nucleotides, nucleosides, nucleotide precursors and salvage intermediates. *RFK* loss substantially depleted nucleotide precursors associated with both purine and pyrimidine biosynthesis (such as carbamoyl aspartate, AICAR, SAICAR and GAR) as well as cytidylate nucleotides and nucleosides (cytosine, cytidine, 2’-deoxycytidine, dCMP and dCDP) (Figure 4C, Table S3). Consistent with the notion that riboflavin perturbation impairs the activity of DHODH, we observed an increased dihydoorotate:orotate ratio and decreased total DHODH protein in two AML cell lines upon *RFK* loss or riboflavin starvation (Figure 4C, 4D and S4A). Importantly, supplementation of aspartate, a key ETC metabolic product necessary for nucleotide biosynthesis, did not rescue cell proliferation upon *RFK* KO, consistent with the notion that DHODH operated downstream of the step that requires aspartate (Figure S4F).

To better understand the molecular basis of these observations, we employed a CRISPR screening approach. Briefly, we designed a metabolism-focused CRISPR-Cas9 library targeting 2,902 metabolic genes with a total of 6,072 sgRNAs including controls (2 sgRNAs per gene) (Figure S4G and Table S4, see Methods). We infected NB4-Cas9 cells with this library and cultured them for 14 days in either Plasmax complete medium or Plasmax lacking riboflavin and compared sgRNA abundance by deep sequencing at endpoint (Figure 4E). Remarkably, loss of most enzymes involved in *de novo* purine biosynthesis and folate metabolism/transport scored as the top mechanisms of resistance to riboflavin starvation, with loss of these genes providing growth advantage in riboflavin deficient medium (Figure 4F, Table S5). As our data showed *RFK* loss specifically depleted cytidylate pools, and loss of most enzymes involved in *de novo* purine biosynthesis confers resistance to exogenous riboflavin deprivation, we hypothesized that nucleotide imbalance between purines and pyrimidines may be a primary contributor for loss of AML proliferative potential upon perturbation of riboflavin metabolism. To functionally confirm our screen results we cultured NB4 cells in Plasmax complete or riboflavin deficient medium and supplemented liquid culture with cocktails of purine (A+G) or pyrimidine (C+T) nucleosides. In accordance with our screen results, only supplementation of pyrimidine nucleotides provided proliferative advantage to cells in riboflavin deficient conditions (Figure 4G). To expand these results, we cultured *RFK* intact or deleted MV4-11 AML cells in Plasmax culture medium supplemented with cocktails of purine or pyrimidine nucleosides. Culture of *RFK* replete cells with either purine or pyrimidine nucleosides decreased proliferation, consistent with previous reporting of antiproliferative effects of supraphysiological levels of exogenous nucleosides^50,51^ (Figure 4H). In *RFK* depleted conditions, addition of purine nucleosides exacerbated the antiproliferative effects of *RFK* loss while supplementation of pyrimidine nucleosides strongly rescued proliferation, supporting our hypothesis that *RFK* loss specifically depletes pyrimidine pools. These results were phenocopied by repeating the experiment in RPMI medium (Figure S4H). As an orthogonal form of validation, it has previously been shown that nucleotide imbalance induces expression of DNA damage markers, such as phosphorylation of histone variant γH2A.X^50,51^. Indeed, we observed that loss of *RFK* induced potent DNA damage that was largely absolved via the supplementation of pyrimidine nucleosides (Figure 4I). Addition of cytidylates alone (cytidine or 2’-deoxycytidine) was insufficient to rescue the antiproliferative effects of *RFK* loss, suggesting maintenance of balanced pyrimidine pools (i.e., upon addition of both cytidylates and thymidylates) is necessary to relieve the effects of riboflavin metabolism perturbation (Figure S4I). Interestingly, the levels of certain cytidylate species can be used as a predictive marker of *RFK* sensitivity; in AML, lower levels of endogenous dCMP and cytidine (although not to a level of statistical significance for cytidine) predict greater dependence on *RFK* in the DepMap dataset (Figure 4J). Collectively, these data illustrate that perturbation of riboflavin metabolism induces a potent nucleotide imbalance response that can be rescued by either attenuating *de novo* purine metabolism or supplementing liquid culture with pyrimidine nucleotides.

### Perturbation of riboflavin metabolism impairs iron-sulfur cluster proteins

To further characterize biological processes which interact with riboflavin metabolism, we examined genes whose loss sensitizes cells to riboflavin starvation in our CRISPR screen. As expected, and consistent with our proteomic data, loss of genes which encode subunits of mitochondrial complex I conferred sensitivity to riboflavin starvation, and deletion of *RFK* or *SLC25A32* (the mitochondrial FAD transporter^52,53^) further sensitized AML cells to riboflavin starvation (Figure 5A and Table S5).

**Figure 5.**
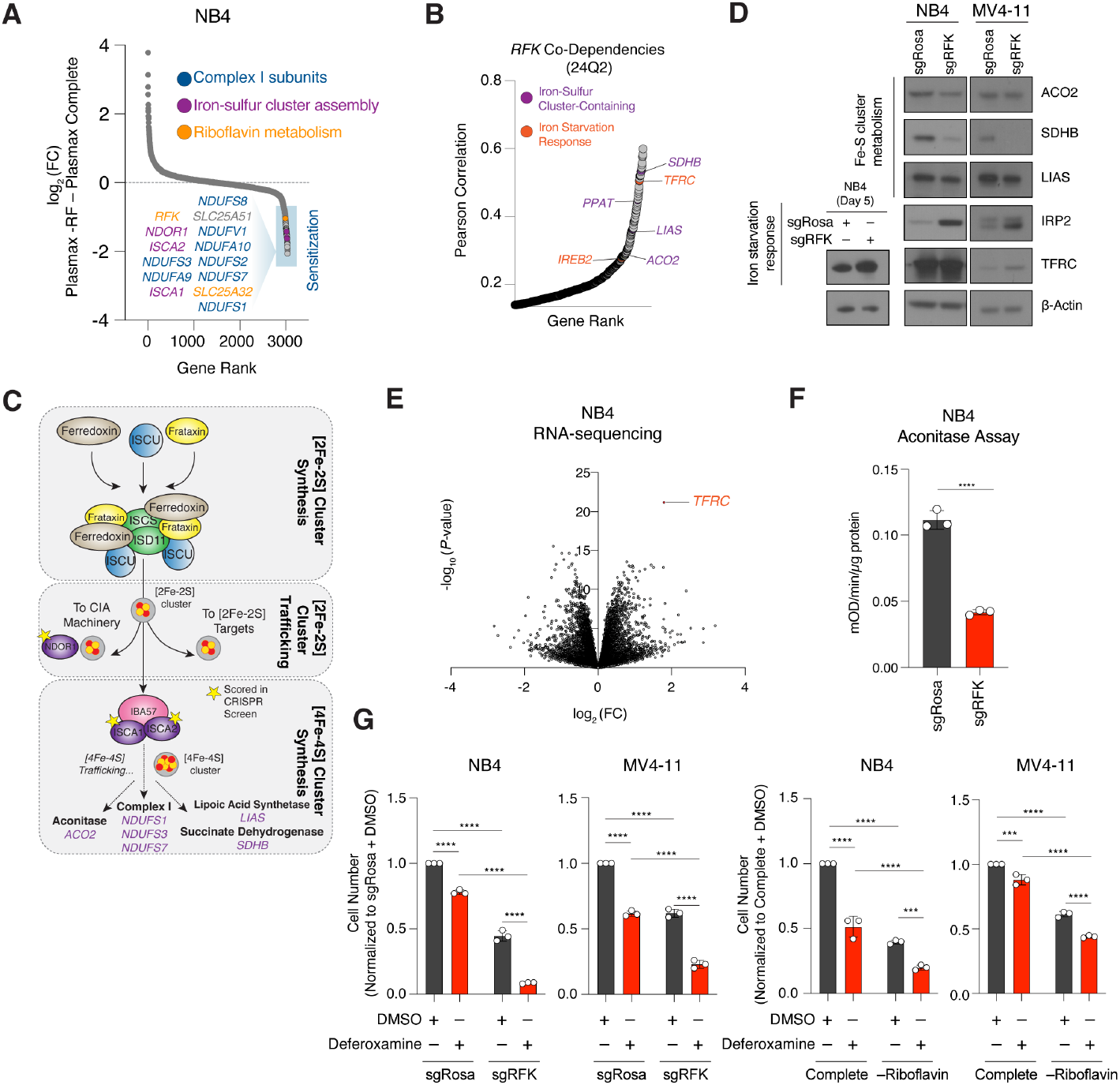
Riboflavin perturbation impairs iron-sulfur cluster dependent proteins. **A**. Dot plot of genes ranked by relative log_2_ enrichment or depletion in Plasmax complete versus riboflavin depleted culture medium as measured by sgRNA abundance. Genes whose abundance decreased conferred sensitivity to riboflavin depletion. Colors show indicated gene families. **B**. Dot plot of genetic co-dependency of genes with *RFK* ranked by Pearson correlation. Data from DepMap 24Q2. Purple shading indicates iron-sulfur cluster containing genes, and orange iron starvation response-associated genes. **C**. Simplified schema of the synthesis of [2Fe-2S] iron-sulfur clusters, their trafficking, and involvement in the synthesis of [4Fe-4S] iron-sulfur clusters. The client proteins of [4Fe-4S] clusters are highlighted, and stars indicate CRISPR screen hits that sensitize cells to death upon riboflavin starvation. **D**. Western immunoblot analysis of indicated proteins in cell lysates isolated from NB4 and MV4-11 cells at day 5 (left blots) and day 9 (right blots) post induction of Rosa or *RFK* sgRNAs. β-Actin served as the loading control. **E**. Volcano plot of relative fold change (log_2_) of differentially regulated genes versus −log_10_ (*P*-values) in NB4 cells upon *RFK* deletion in RPMI medium at day 7. **F**. Aconitase activity (mOD/min/μg of protein) in whole-cell lysates of *RFK* deleted NB4 cells at day 9. mOD, milli optical density. Data representative of *n*=3 biological replicates. **G**. Cell number of NB4 and MV4-11 cells at day 7 post induction of Rosa or *RFK* sgRNAs, or day 7 post riboflavin starvation, treated with vehicle (DMSO, dimethyl sulfoxide), or the iron chelator deferoxamine (DFO) for 3 days. Cell number normalized to sgRosa + DMSO, or Plasmax Complete medium + DMSO. Data representative of *n*=3 biological replicates. Data are presented as the mean ± SD. *** *P* < 0.001, \*\*\*\**P* < 0.0001 by unpaired two-tailed Student’s *t*-test (F) and ordinary two-way ANOVA with uncorrected Fisher’s LSD test (G).

A recurrent biological signature identified in our proteomic dataset is a relationship between riboflavin metabolism and proteins that contain an iron-sulfur (Fe-S) cluster. Namely, loss of *RFK* reduced the abundance of a number of proteins requiring Fe-S clusters but not FMN/FAD as cofactors (Figure 3B and 3D). Moreover, several canonical enzymes that require Fe-S clusters are genetic co-dependencies with *RFK* (such as the iron-sulfur binding subunit of succinate dehydrogenase (*SDHB*), aconitase 2 (*ACO2*), and lipoic acid synthetase (*LIAS*), along with genes which control iron starvation responses, such as *IREB2* and the transferrin receptor (*TFRC*) (Figure 5B)). Fe-S clusters are ancient inorganic co-factors that have been highly conserved throughout evolution and are involved in electron transfer and catalysis reactions^54^. While the chemical structure of Fe-S clusters is relatively simple, their biosynthetic pathways are highly complex, with over 30 proteins directly involved in the synthesis and assembly of these cofactors^55^. The simplest form of a Fe-S cluster is the rhombic [2Fe-2S] cluster, which contains two iron atoms connected by two sulfide ions; another is the cuboid [4Fe-4S] cluster, which contains four iron and four sulfide ions. Both cluster types can adopt various oxidation states and bind distinct partner proteins^56^. Remarkably, in our CRISPR screen, knockout of Iron-Sulfur Cluster Assembly 1 and 2 (*ISCA1* and *ISCA2*) scored as strongly sensitizing to riboflavin deprivation (Figure 5A and 5C). After synthesis of a [2Fe-2S] cluster, ISCA1 and ISCA2 form a heterodimeric complex and are involved in its mitochondrial conversion to the [4Fe-4S] cluster, a process that is FMN and FAD-independent^57,58^. Previous work demonstrated that loss of *ISCA1* or *ISCA2* ablates mitochondrial structural integrity and results in highly specific loss of function of [4Fe-4S]-dependent proteins, including mitochondrial complex I and II, LIAS and aconitases^57^ – all genetic co-dependencies of *RFK* (Figure 5B). Our screen did not identify any genes involved in the early core synthesis of Fe-S clusters, but did find that loss of *NDOR1*, a gene involved in assembly of cytosolic Fe-S clusters also sensitizes to riboflavin loss (Figure 5A). NDOR1 complexes with CIAPIN1 and oxidizes NADPH via its FMN and FAD cofactors and delivers electrons to cytosolic Fe-S scaffolds^59,60^. These results suggest a model whereby the phenotypes observed in *RFK* KO or riboflavin starved cells could in part result from an Fe-S cluster deficiency, and riboflavin metabolism intersects with Fe-S cluster metabolism specifically at the level of [4Fe-4S]-dependent proteins. Indeed, *RFK* KO in two AML cell lines reduced the protein abundance of several Fe-S dependent proteins, including ACO2, which mediates the conversion of citrate to isocitrate in the TCA cycle, SDHB, and LIAS (Figure 5D). This was accompanied by a well-characterized Fe-S cluster iron starvation response whereby IRP2, an iron deficiency sensor, stabilizes the mRNA transcripts of the transferrin receptor (TFRC) to import extracellular iron. These results were largely phenocopied in the context of exogenous riboflavin starvation with the exception of ACO2, which appeared to remain stable (Figure S5A). Strikingly, in an RNA-sequencing dataset generated in *RFK* KO AML cells, *TFRC* was the most highly up-regulated transcript at day 7 (Figure 5E, S5B and S5C, Table S6). Additionally, *RFK* KO cells showed functionally impaired whole-cell aconitase activity, a class of enzymes which require Fe-S clusters (Figure 5F). Enhanced sensitivity to chelation of intracellular iron with the iron chelator deferoxamine was also observed in *RFK* KO cells, as well as in riboflavin starvation conditions (Figure 5G). Finally, similar to the levels of certain cytidylate species, aconitate, an intermediate in the conversion of citrate to isocitrate in the TCA cycle, is among the highest scoring predictive markers of *RFK* sensitivity. In AML, lower levels of aconitate predict greater dependence on *RFK* in the DepMap dataset, albeit not to statistical significance (Figure S5D). Thus, these results support the contention that *RFK* loss impairs the activity and stability of proteins containing iron–sulfur cluster co-factors.

### Riboflavin metabolism perturbation synergizes with BCL-2 inhibition

Finally, we sought to identify modalities by which we could therapeutically exploit riboflavin metabolism in AML. We took advantage of previous reporting that inhibition of the ETC induces mitochondrial dysfunction by priming cells for apoptosis via activation of pro-apoptotic BH3-only proteins such as BIM and NOXA in an ATF4-dependent manner and shifting pro-survival cues to the anti-apoptotic protein BCL-2^61^. The BCL-2 inhibitor venetoclax is combined with hypomethylating agents to treat patients with AML unfit for intensive chemotherapy^62^. We thus reasoned that the collapse in oxidative phosphorylation observed upon perturbing riboflavin metabolism might sensitize AMLs to venetoclax. To test this hypothesis, we used genetic, exogenous depletion and small molecule approaches (Figure 6A). As predicted, genetic KO of *RFK* sensitized multiple AML cell lines to venetoclax, including PDX16-01, a model which we have observed to be highly resistant to single-agent venetoclax treatment^24^ (Figure 6B). As an orthogonal approach, we cultured AML cells in Plasmax complete medium or Plasmax lacking riboflavin and similarly observed increased sensitivity to venetoclax upon exogenous riboflavin withdrawal (Figure 6C). We were unable to culture our PDX cells *in vitro* in Plasmax to test venetoclax sensitivity upon exogenous depletion of riboflavin.

**Figure 6.**
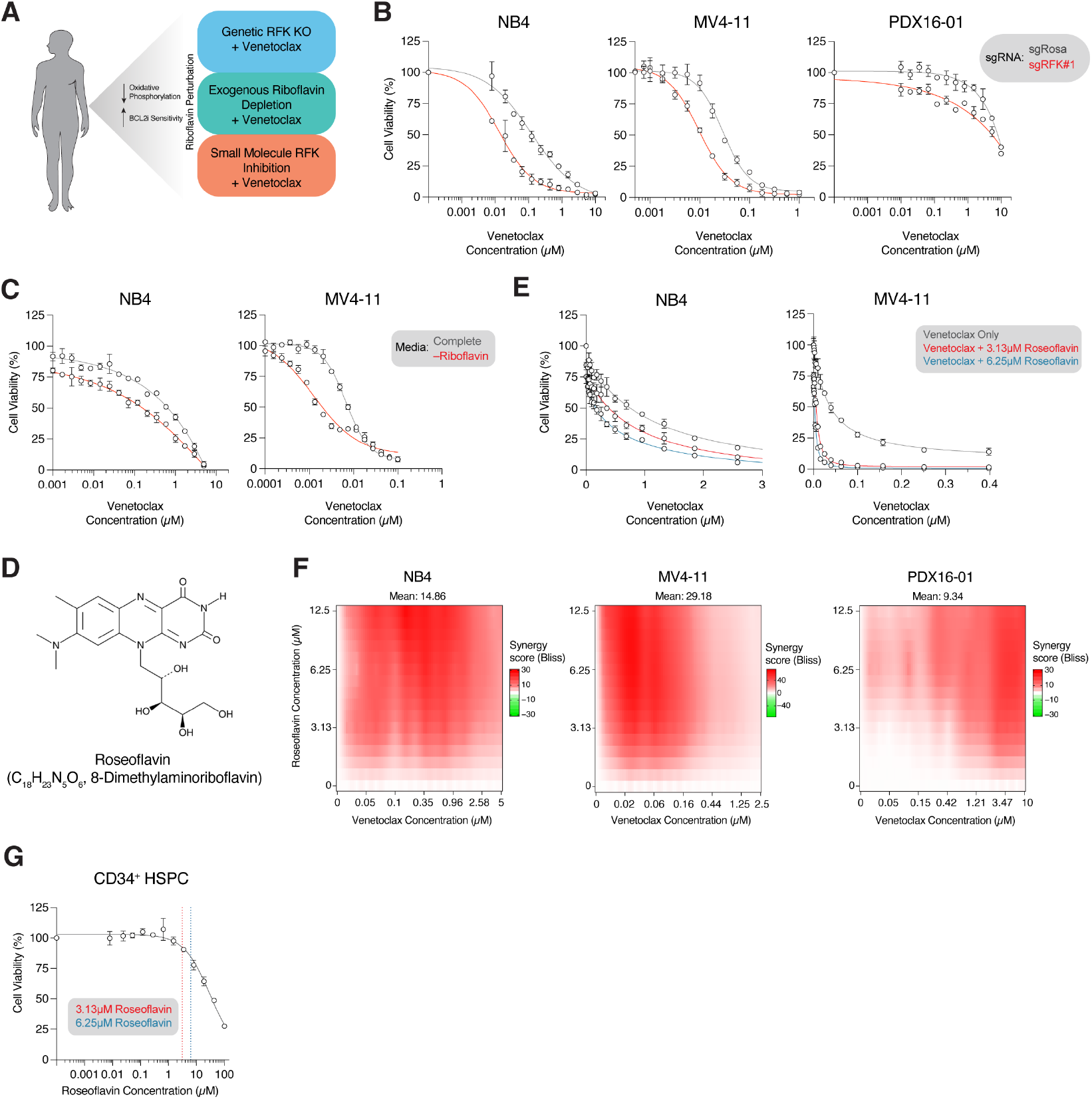
Perturbing riboflavin metabolism synergizes with BCL-2 inhibition. **A**. Schema of RFK perturbation via genetic, exogenous depletion and small molecule approaches in combination with the BCL-2 inhibitor venetoclax. **B**. Relative viability of doxycycline-inducible sgRNA-expressing NB4, MV4-11 and PDX16-01 cells targeting Rosa or *RFK* treated with venetoclax for 72 hours. Cells were treated with doxycycline for 4 days prior to venetoclax treatment to induce gene knockout. **C**. Relative viability of NB4 and MV4-11 cells cultured in Plasmax complete or riboflavin deficient medium and treated with venetoclax for 72 hours. Cells were pre-conditioned in appropriate medium for 4 days prior to venetoclax treatment. **D**. Chemical structure of roseoflavin, 8-dimethylaminoriboflavin. **E**. Relative viability of NB4 or MV4-11 cells treated with venetoclax alone, or venetoclax and two fixed concentrations of roseoflavin (3.13 μM, blue; or 6.25 μM, red) for 72 hours. **F**. NB4, MV4-11 and PDX16-01 cells were treated with escalating concentrations of venetoclax and roseoflavin for 72 hours to determine viability effects. The presence of treatment synergy was determined using SynergyFinder and the Bliss synergy index and is denoted as regions of red in the graphs. The mean of three biological replicates was used for each data point. **G**. Relative viability of CD34^+^ HSPC cells treated with escalating concentrations of roseoflavin for 72 hours. Concentrations used to sensitize NB4 and MV4-11 cells in E. are highlighted (3.13 μM, blue; or 6.25 μM, red). Data are presented as the mean ± SD.

While the above experiments provide proof of concept that targeting RFK enhances venetoclax sensitivity in AML, it is likely that absolute RFK loss-of-function harbors significant off-tumor hematopoietic toxicities compared to the transient inhibition afforded by small molecule inhibitors. Indeed, CRISPR-based deletion of *RFK* in healthy donor derived human CD34^+^ HSPCs uniformly impaired colony forming capacity (Figure S6A).

In the absence of bona fide RFK inhibitors, we turned our attention to roseoflavin, a *Streptomyces davawensis*-derived antimetabolite analog of FMN and riboflavin that has previously been employed as a putative antibacterial agent^63^ and has been shown to bind and be metabolized by human RFK^64^ (Figure 6D). Consistent with our genetic and exogenous withdrawal approaches, the addition of roseoflavin at two fixed low-micromolar concentrations significantly sensitized AML cell lines to venetoclax (Figure 6E). Indeed, combinatorial dose-escalation of roseoflavin and venetoclax proved strongly synergistic in multiple AML cell lines and PDX16-01 as demonstrated by the Bliss synergy index (Figure 6F and S6B). Importantly, roseoflavin treatment killed healthy CD34^+^ HSPCs at high-micromolar concentrations but only sparingly at the concentrations used to sensitize AMLs to venetoclax, suggesting a potential therapeutic window (Figure 6G). Thus, these experiments provide a potential roadmap for future therapeutic approaches harnessing riboflavin metabolism as a mechanism to sensitize AMLs to clinically used therapeutics.

## Discussion

In this study, we identified riboflavin metabolism as a vulnerability in AML and its essential contribution to mitochondrial respiration, nucleotide biosynthesis, and iron-sulfur cluster metabolism in this disease. Consistent with a wide body of previous literature, we confirmed the essentiality of FMN and FAD in the maintenance of oxidative phosphorylation and showed that loss of these cofactors specifically results in destabilization and attenuated assembly of mature mitochondrial complex I and II. Previous work has shown that exogenous depletion of riboflavin or genetic KO of the mitochondrial FAD transporter *SLC25A32* in mouse fibroblasts or 143B cells impairs complex I and II assembly^42^. Similarly, riboflavin deficiency in B16 mouse melanoma cells induces broad destabilization and proteolytic turnover of flavin-dependent enzymes^65^. Our work supports and extends these observations in the context of leukemia and demonstrates how flavin deficiency-induced protein destabilization, particularly regarding complex I and II, can be employed as a strategy for inducing sensitivity to targeted therapies such as BCL-2 inhibitors. Given the practical difficulty in enforcing riboflavin deficient diets in patients with cancer, our work employed roseoflavin, a small-molecule antimetabolite of riboflavin and FMN, and demonstrated synergy with venetoclax and a potential therapeutic window when comparing response in AML to normal CD34^+^ HSPCs. Investment in developing *in vivo*-optimized riboflavin anti-metabolites/RFK inhibitors is warranted. Studies in the 1960s trialed antimetabolites of riboflavin such as galactoflavin in humans and documented mild clinical signs of riboflavin deficiency such as sore throat, tongue changes and angular stomatitis at 3 weeks post initiation of therapy. More serious effects such as peripheral neuropathy were observed at 5 weeks and partially improved upon cessation of treatment and riboflavin add-back^66^. Thus, these proof-of-concept studies suggest that transient inhibition of RFK may induce tolerable and/or manageable side effects while offering a window of synergistic opportunity with venetoclax. Rescue with high-dose systemic riboflavin supplementation after treatment, akin to leucovorin supplied with methotrexate, may further reduce side effects^67^.

Another downstream consequence of perturbing riboflavin metabolism is the specific depletion of pyrimidines and induction of a nucleotide imbalance DNA damage response. An attractive explanation of this imbalance is reduced activity of DHODH, an FMN-dependent protein that is involved in the second step of pyrimidine metabolism and which has itself been explored as a potential AML therapeutic target^49^, though other mechanisms may be at play. The reduced abundance of a number of DNA repair enzymes, such as NTHL1 and DDX11 upon *RFK* loss, may also contribute to the DNA damage phenotype we observed. Additionally, it has been shown in HepG2 cells that riboflavin starvation *in vitro* can induce oxidative stress via reactive oxygen species production and increased DNA strand breaks (ostensibly due to impaired activity of FAD-dependent glutathione reductase)^68^. Given that many chemotherapeutic agents induce DNA damage, a logical extension of this work would be to determine whether these agents are also potentiated by perturbing riboflavin metabolism.

Perhaps the most intriguing observation in this study is the recurrent intersection of riboflavin and Fe-S cluster metabolism. The relationship between iron and riboflavin has long been appreciated. In bacteria, physiological availability of iron affects the synthesis of riboflavin, as prokaryotes maintain the capacity for *de novo* synthesis. Indeed, iron starvation or deletion of *Fur*, the primary regulator of iron homeostasis in bacteria, increases transcription of the riboflavin synthesis operon in *Clostridium acetobutylicum*^69^ and iron availability and *Fur* act as negative regulators of riboflavin biosynthetic operons in other bacteria^70,71^. In humans, clinical riboflavin deficiency impairs iron handling and has been observed to enhance iron absorption and decrease intracellular iron mobilization; many patients are also anemic^72,73^. In the absence of riboflavin biosynthesis in humans, our data argues that these physiological responses can, at least partially, be explained as a Fe-S cluster starvation response induced upon interruption of riboflavin metabolism. Indeed, our data support a model whereby a key, previously unappreciated role for riboflavin is mediation of Fe-S cluster biosynthesis. Recent work has shown that perturbation of mitochondrial glutathione (GSH) induces remarkably similar physiological responses to those we observed (particularly analogous iron starvation responses), highlighting the complexity of mitochondrial Fe-S biogenesis and the contribution of previously non-canonical metabolites in their synthesis^74^.

Our CRISPR screening data provides intriguing clues as to how riboflavin may interact with Fe-S metabolism. *ISCA1* and *ISCA2* are evolutionarily conserved genes that play essential roles in the assembly and maturation of [4Fe-4S] iron-sulfur clusters and participate in their transfer to client proteins in the mitrochondria^57^. Indeed, loss of *ISCA1* and *ISCA2* specifically reduces the activity of [4Fe-4S]-dependent proteins while proteins binding [2Fe-2S] clusters are unaffected^57,75^. The highly specific depletion of *ISCA1* and *ISCA2*, independent of any other genes involved in core Fe-S synthesis, suggests that RFK likely acts in a parallel or compensatory pathway to [4Fe-4S] cluster assembly or utilization. Given the known roles of riboflavin in maintaining cellular redox homeostasis (e.g., glutathione-disulfide reductase requires a FAD cofactor), this mechanism may center on maintaining reduced glutathione pools to enable iron chaperone activity^76^, potentially for the Fe-S biosynthesis pathway, or by protecting [4Fe-4S] clusters from oxidative damage; for instance, superoxide accumulation is known to rapidly inactivate [4Fe-4S] clusters, and superoxide dismutases (SODs) are sensitive to riboflavin abundance^77^.

## Limitations of This Study

Though this manuscript identified that hematological malignancies have heightened dependency on *RFK*, it is unclear why a large proportion of solid tumor cell lines appear to have dispensable requirement for this enzyme. This may be a result of lineage-specific compensatory/bypass mechanisms which we have not explored in our study. Additionally, while we identify impaired DHODH activity as the likely primary cause for pyrimidine depletion upon perturbation of riboflavin metabolism, the mechanisms of specific cytidylate depletion are unclear. It is important to note, however, that cytidylate add-back alone is insufficient to rescue proliferative defects caused by riboflavin metabolism perturbation, and a cocktail of complete pyrimidines is required to reverse this phenotype. Finally, though we present substantial data linking riboflavin metabolism to [4Fe-4S] Fe-S cluster dependent enzymes, the exact mechanistic basis of this connection is not clear; further work is required to elucidate these mechanisms.

## Resource Availability

### Lead Contact

Further information and requests for resources should be directed to and will be fulfilled by the lead contact, Kimberly Stegmaier.

### Materials Availability

The metabolic CRISPR-Cas9 library generated in this study will be deposited to Addgene and released upon publication of this manuscript. The Broad Institute Clone Pool (CP) identifier for this library is: CP1957.

### Data and code availability

The proteomics data generated by RFK deletion and exogenous riboflavin deprivation will be deposited to the Proteomics Identifications Database (PRIDE) and released upon publication of this manuscript.

Raw RNA-sequencing data generated upon RFK KO will be deposited to the NCBI Gene Expression Omnibus and released upon publication of this manuscript.

Metabolomics and CRISPR screen data is available in the Supplementary Tables.

Original source data and any additional required information are available from the lead contact upon reasonable request.

## Acknowledgements

S. Bjelosevic is a Fellow of and was supported by the Leukemia & Lymphoma Society (LLS). R. Fauth was supported by a Boehringer Ingelheim Fonds MD Fellowship. B.T. Do was supported by NHLBI (F30HL156404) and NIGMS (T32GM007753). T.D. Jackson is supported by a Cancer Research Institute Irvington Postdoctoral Fellowship (CRI5298). C. Schneider reports salary support from the German Research Foundation (Deutsche Forschungsgemeinschaft, SCHN 1622/1-1) and Helen Gurley Brown Foundation. S. Lin was a Fellow of the Leukemia and Lymphoma Society and was supported by NCI R00CA263161. M.G. Vander Heiden acknowledges support from the Ludwig Center at MIT, the MIT Center for Precision Cancer Medicine, a Faculty Scholar award from HHMI; a Koch Institute/DFHCC Bridge Grant and the NCI (P30CA14051; R35CA242379). K. Stegmaier was supported by funding from The Selig Family Fund for Pediatric Cancer Research, the Children’s Leukemia Research Association (CLRA), the National Cancer Institute R35 CA283977, and the St. Jude Children’s Research Hospital Transcription Collaborative Research Program. We acknowledge and thank the Genetic Perturbation Platform (GPP) and DepMap/PedDep Team at the Broad Institute of MIT and Harvard, The Thermo Fisher Center for Multiplexed Proteomics at Harvard Medical School, the Whitehead Institute Metabolomics Core, Dana-Farber Cancer Institute Metabolomics Facility, and Dana-Farber Cancer Institute Animal Resources Facility (ARF) for invaluable contributions to this work. We thank members of the Stegmaier, Vander Heiden and Puigserver laboratories for their constructive feedback and expertise.

## Author Contributions

S.B. and K.S. conceived the project, analyzed and interpreted the data and wrote the manuscript with input from all co-authors. K.S. secured funding for the study. S.B., R.F. and L.A.M. conducted most of the experiments. B.T.D, R.E. and M.B.M. conducted metabolomic analysis and provided expertise. G.A. and G.D. performed CRISPR screen and RNA-sequencing computational analysis. L.A.M., A.T.B., A.T. and S.S. conducted mouse experiments. B.R. performed metabolomics processing and analysis and provided expertise. T.D.J. performed BN/SDS-PAGE and analysis with expertise and funding from P.P. C.S. and N.B. provided experimental expertise and conceptual guidance. S.L. generated Cas9-competent PDX models, previously performed in vivo CRISPR screens and provided expertise. D.E.R. generated the metabolic CRISPR screen library at the Broad Institute Genomic Perturbation Platform (GPP) and provided critical technical and conceptual expertise. M.G.V.H. provided critical expertise and funding for metabolomics experiments.

## Supplemental Information Titles and Legends

Document S1. Figures S1–S6 (embedded in this document)

Table S1. Proteomics data related to Figure 3 and S3

Table S2. Metabolite profiling data related to Figure 4 and S4

Table S3. Metabolite profiling related to Figure 4 and S4

Table S4. Metabolism-focused CRISPR library

Table S5. CRISPR screen data related to Figure 4 and S4; Figure 5 and S5

Table S6. RNA-sequencing data related to Figure 5 and S5 Table S7. CRISPR sgRNA and cDNA sequences

## Materials and Methods

### Experimental Models and Study Participant Details Cell Lines and PDX Samples

All commercially available cell lines were purchased from either ATCC (MV4-11, U937, SKNAS and HEK-293T) or DSMZ (NB4, MOLM-13, OCI-AML2, OCI-AML3 and EW8). MV4-11, U937, NB4, MOLM-13, OCI-AML2, OCI-AML3 and EW8 were cultured at baseline in Roswell Park Memorial Institute 1640 (RPMI-1640), supplemented with 10% fetal bovine serum (FBS) and 1% penicillin-streptomycin (P/S). SKNAS and HEK-293T cells were cultured at baseline in Dulbecco’s Modified Eagle Medium (DMEM), supplemented with 10% fetal bovine serum (FBS) and 1% penicillin-streptomycin (P/S). Cell line identity was confirmed via short tandem repeat (STR) profiling by LabCorp at the start of the experiment. Mycoplasma contamination was routinely tested for using the MycoAlert Mycoplasma Detection Kit. All cell lines were cultured at 37°C and at 5% atmospheric CO_2_. Cells were maintained and most experiments performed in base medium, unless otherwise noted in the manuscript. For experiments in which a transition to physiological medium (Plasmax) was performed, cells were equilibrated for at least 48 hours in Plasmax complete preculture prior to commencing experiments.

Derivation and establishment of PDX models was described previously^24^. For short-term *in vitro* culture, PDX16-01 and PDX17-14 were maintained in Iscove’s Modified Dulbecco’s Medium (IMDM), supplemented with 20% fetal bovine serum (FBS), 1% penicillin-streptomycin (P/S) and 10 ng/mL of human SCF, TPO, FLT3L, IL3, and IL6. A summary of the genetics of these two models is provided below:

**PDX16-01**, Relapsed AML, CALM-AF10, complex karyotype, NF1, PHF6, TP53 mutant

**PDX17-14**, Secondary AML, relapsed MLL-AF10, complex karyotype, KRAS mutant

### Primary Human CD34+ HSPC Culture

CD34^+^ hematopoietic stem/progenitor cells (HSPCs) were purchased from Lonza, and each vial contained 1 × 10^6^ cells with greater than 90% CD34+ purity. Donors were deidentified and negative for standard virology screening, and informed consent was provided. Vials from Donor ID# 47554, 48310 and 50616 were obtained for this study. Cells were thawed and stimulated in StemSpan II medium supplemented with 100 ng/mL human SCF, TPO and FLT3L, and 10 ng/mL IL3 and IL6 for 48 hours prior to transition to appropriate target cell medium.

### Animal Models

All animal experiments were performed in accordance with the approved protocols from the Dana-Farber Cancer Institute Institutional Animal Care and Use Committee (IACUC) and performed in accordance with protocols approved under protocol #15-029. NOD-*scid* IL2Rg^null^ (NSG) female mice (6-8 weeks old) were obtained from the Jackson Laboratory and used for diet manipulation and xenograft experiments. Mice were maintained in ventilated cages and fed with sterile food and water at the Longwood Center Animal Resource Facility. Mice were housed at ambient temperature and humidity (18-23ºC, 40-60% humidity) with a 12-hour light-dark cycle and co-housed with littermates with *ad libitum* access to water and appropriate diet. All experimental subjects were age-matched, numbered and randomly assigned. Data is reported as *n* equaling biologically independent animals. Statistical methods were not used to calculate sample size.

### Generation of CRISPR-Cas9 Cell Lines and CRISPR-Mediated Gene Knockout

To generate Cas9 competent AML and the EW8 cell lines, we stably expressed the FU-Cas9-mCherry vector in parental cells via lentiviral transduction. Cells were then FACS sorted as necessary using a BD FACSAria III on mCherry expression until a homogenous population was established. Cas9-competent SKNAS cells were generated by transducing cells with the LentiCRISPRV2-Blast plasmid and selecting and maintaining cells in culture medium supplemented with 10 μg/mL blasticidin. Cas9 activity was confirmed using the Broad Institute’s Cas9 eGFP Reporter Assay (using pXPR-047 and pRDA-221 plasmids). Cells were STR profiled to ensure post engineering identity. To achieve doxycycline-inducible CRISPR-mediated deletion of target genes, we cloned annealed oligonucleotide sequences targeting the gene of interest into the FgH1t-UTG-GFP sgRNA scaffold plasmid and transduced it into target cells. Cells were subsequently FACS sorted for the double-positive GFP/mCherry population (i.e., sgRNA-GFP and Cas9-mCherry). Induction of sgRNAs was achieved via supplementation of 1 μg/mL doxycycline into cell culture for 96 hours.

To generate lentivirus for the above transductions, 6 × 10^6^ HEK-293T cells were seeded in 10 cm dish 24 hours prior to transfection, and media refreshed on the day of transfection. 3 μg of psPAX2 lentiviral packaging plasmid, 1.5 μg of pMD2.G VSV-G envelope plasmid, along with 2 μg of target plasmid were transfected into cells using TransIT reagent. 24 hours post transfection media was refreshed, and viral supernatant collected 48 hours post transfection by filtration through a 0.45 μm cell filter. To transduce target cells, 3 × 10^6^ cells were seeded in a well of a 12-well plate, and 0.1 – 1 mL of viral supernatant added to each well in addition to 6-8 μg/mL of polybrene reagent. Cells were then spinfected at 900 *g* for 2 hours at 30ºC to enhance infection efficiency.

### Plasmax Synthesis and Vitamin Depletion Screens

Detailed protocols establishing Plasmax have been previously described^20^. Briefly, we prepared stock solution aliquots of each of the key Plasmax constituents: 100X amino acid stock (solution #1 – adjusted to pH 1.0 to increase solubility); 100X miscellaneous polar metabolites (solution #2); 1000X trace metals/minerals (solution #3); 500X urate (solution #4 – adjusted to pH 13.30 to increase solubility); and 100X vitamin stock (solution #5, supplemented with ascorbate and vitamin B_12_). Each master stock was filtered with a 0.22 μm vacuum filtration filter and stored at −80ºC. To assemble a bottle of Plasmax complete medium, solution #1, 2, 3, 4 and 5 was added to 500 mL of commercially purchased Earle’s Balanced Salt Solution (EBSS), along with L-glutamine, sodium pyruvate, and 5% dialyzed fetal bovine serum (dFBS). Each bottle was pH equilibrated at 5% CO2, and we confirmed a final pH of ~7.2 per bottle. For metabolite depletion experiments, we synthesized stock solutions omitting the metabolite of interest; for example, for riboflavin depletion studies, we made up solution #5 omitting riboflavin but maintaining all other metabolites at appropriate concentrations. Plasmax was then synthesized as above.

To perform the vitamin depletion screen, 0.2 × 10^6^ NB4 or MOLM-13 cells in 5 mL of medium were seeded in each well of a 6-well plate in duplicate. As an untreated control, cells were cultured in Plasmax complete. As a positive antiproliferative/differentiation control, cells were cultured in Plasmax complete treated with 1 μM of all-*trans* retinoic acid (ATRA). For all other conditions, cells were cultured in Plasmax complete omitting solution #5; vitamins were manually supplemented at appropriate concentrations to each well omitting the vitamin of interest. To test for proliferation effects of vitamin depletion, after 72 hours, 300 μL of thoroughly homogenized cell suspension was harvested from each well, and 30 μL of CountBright Absolute Counting Beads were spiked into the suspension. 2000 events were acquired via flow cytometry on a BD FACSCelesta, with end gating set on bead events and co-capturing live cells. To test for changes in cell state/differentiation upon vitamin depletion, 200 μL of cell suspension was concurrently harvested from each well, spun down at 400 *g* for 4 minutes, and media aspirated. Cells were resuspended in phosphate buffered saline (PBS), 0.6 μL of CD11b-APC antibody per 500,000 cells, and stained at room temperature and in the dark for 15 minutes. Cells were then washed twice in PBS and processed on the flow cytometer. Remaining cells in each well were spun down and resuspended in appropriate fresh Plasmax medium as above for 72 hours. At day 6, the harvest procedure was repeated. The experiment was repeated twice, and four pooled replicates were used for each data point in the screen across the two repeats. Data was analyzed via FlowJo software and acquired bead count/live cell count used to calculate cells/μL of culture. Cell counts were normalized to the Plasmax complete condition.

### Annexin V Apoptosis and Differentiation Assays

To quantify induction of apoptosis upon genetic loss of *RFK*, doxycycline-inducible sgRNAs were induced with 1 μg/mL doxycycline, and cells cultured for the specified time period. To quantify induction of apoptosis upon exogenous withdrawal of riboflavin, cells were pre-cultured in Plasmax complete for 48 hours, and then in Plasmax complete or riboflavin deficient medium for the specified time period. Cell suspension was harvested and cells washed in PBS prior to re-suspension in Annexin V Binding Buffer, containing 1:100 diluted Annexin V-APC antibody. Samples were vortexed and stained in the dark at room temperature for 15 minutes prior to analysis on a BD FACSCelesta.

For assessment of differentiation changes in AML cells upon loss of *RFK*, cells were induced as above, harvested and washed at endpoint. Cells were re-suspended in PBS containing 0.6 μL of CD11b-APC antibody per 500,000 cells, or 0.6 μL of appropriate isotype control (IgG2b, κ). Cells were stained at room temperature and in the dark for 15 minutes, washed twice, and analyzed on a BD FACSCelesta.

### *In Vitro* Competitive Proliferation Assays

For proliferative competition experiments, Cas9-competent cells were transduced with doxycycline-inducible sgRNAs tagged with GFP targeting the gene of interest at a multiplicity of infection (MOI) of ~0.8 (i.e., ~80% of cells are GFP positive post infection, and ~20% of the culture is untransduced). Cells were allowed to equilibrate for 72 hours. Cells were then seeded in a 12-well plate at a density of 0.2 × 10^6^ cells/well in triplicate, and a baseline timepoint 0 sample captured via flow cytometry on a BD FACSCelesta to determine starting GFP-positivity. Doxycycline was added to cultures to initiate sgRNA activity at 1 μg/mL. Every 72-96 hours, GFP-positivity was assessed via flow cytometry and the change in the fraction of cells expressing GFP was monitored and cells passaged as required. Where Plasmax culture medium was utilized, cell media was refreshed every 72 hours to prevent nutrient exhaustion. In all experiments, FlowJo software was used for analysis.

### RFK cDNA Overexpression Experiments

To overexpress wild type (WT) *RFK* cDNA in target cells, the coding sequence (CDS) of *RFK* was obtained from NCBI Nucleotide, and sgRNA targeting sequences silently mutated to render resistance to CRISPR-Cas9 mediated deletion. *attB1* and *attB2* overhang sequences were added in-frame with the *RFK* CDS, and a gBlock double-stranded gene fragment was obtained from IDT and subsequently cloned into the pDONR-221 donor vector using the BP Clonase (Gateway Cloning). Successful insertion was validated by whole-plasmid next generation sequencing (NGS). The *RFK*-WT sequence was then cloned into the pLEX_307 destination vector using the LR Clonase, whereby *RFK*-WT expression was downstream of an EF1α promotor and in-frame with a V5 tag along with a puromycin resistance cassette. Successful cloning was validated by whole-plasmid NGS.

NB4 or PDX16-01 cells already stably transduced with doxycycline-inducible sgRNAs targeting endogenous *RFK* were then transduced with the pLEX_307-RFK-WT vector as described above and simultaneously treated with 1 μg/mL of doxycycline to deplete endogenous *RFK*. 24 hours post transduction, cells were selected with 1 μg/mL of puromycin for 72 hours. Post selection, 0.15 × 10^6^ (NB4) or 0.25 × 10^6^ (PDX16-01) cells/well were seeded in a 12-well plate in triplicate, along with appropriate sgRosa or sgRFK alone controls. Cells were counted with CountBright Absolute Counting Beads and flow cytometry as described above. Cells were passaged twice (NB4) or five times (PDX16-01) during the course of the experiment, and cumulative cell counts utilized to determine proliferation rate.

### CD34^+^ HSPC Metabolite Depletion and Roseoflavin Treatment Assays

CD34^+^ HSPCs were thawed and stimulated as described above and transitioned to Plasmax complete, Plasmax minus isoleucine or Plasmax minus riboflavin (all supplemented with the same cocktail of cytokines) as required. For proliferation studies upon amino acid/riboflavin depletion, cells were seeded at 0.2 × 10^6^ cells/well of a suspension-culture coated 12-well plate in triplicate and cell numbers determined via CountBright Absolute Counting Beads and flow cytometry at day 6 and day 14 as described above. The experiment was repeated using material from two separate donors.

For sensitivity to roseoflavin, thawed CD34^+^ HSPCs were seeded at 7,500 cells/well of a 384-well white-bottom plate in 50 μL of Plasmax complete medium supplemented with cytokines and treated with DMSO or escalating concentrations of roseoflavin (0.008 μM to 100 μM) with three replicates for each concentration using a HP D300e Digital Dispenser. After 72 hours, 10 μL of CellTiter-Glo (CTG) was added to each well with gentle agitation for 15 minutes prior to detection of luminescence with a ClarioStar PLUS plate reader. For analysis, background well readings were subtracted from each well, and cell viability expressed as a ratio of signal for each roseoflavin-treated well relative to DMSO. The experiment was repeated using material from two separate donors.

### *RFK* Deletion via Electroporation and Colony Formation Assays

For deletion of *RFK* in CD34^+^ HSPCs, cells were thawed and stimulated for 48 hours in StemSpan II medium and cytokines as above. 6 μg of purified Cas9 nuclease was combined with 100 pmol chemically modified synthetic sgRNA in P3 buffer from the Lonza P3 Primary Cell 4D-Nucleofector X Kit S for 10 minutes at room temperature to form ribonucleoproteins (RNPs). Target CD34^+^ HSPC cells were washed and resuspended at 0.25 × 10^6^ cells per reaction in P3 buffer and combined with the RNP mixture. 25 μL of cell/RNP suspension was transferred into one well of a 16-well Nucleocuvette Strip and Program DZ-100 was used to perform the electroporation on a Lonza 4D Nucleofector Unit. Cells were transferred into a 24-well plate containing pre-warmed medium and were allowed to recover and expand for 48 hours.

For colony formation assays, nucleofected cells were counted and resuspended in MethoCult Express methyl-cellulose media at a concentration of 2,000 cells/mL, with remaining cells from each condition cultured in a 24-well plate in StemSpan II medium with cytokines for confirmation of *RFK* editing (see below). Cells were vigorously vortexed to ensure homogenous suspension and allowed to rest until the solution was bubble-free. 1 mL of cell suspension was then plated in triplicate on 35 mm cell culture plates per condition and incubated at 37ºC at 5% CO2 for 15 days. At endpoint, a 1:1 mixture of PBS and 3-(4,5-Dimethylthiazol-2-yl)-2,5-diphenyltetrazolium bromide (MTT) was prepared, and 200 μL of MTT stain solution was added to each 35 mm cell culture plate for 4 hours at 37ºC. Images were acquired of each plate using a GE ImageQuant LAS 4000, and colony number scored using the “Colony Counting” function in GE ImageQuant TL software.

To verify efficient on-target editing of *RFK* in CD34^+^ HSPCs, we performed Sanger sequencing coupled with Tracking of Indels by DEcomposition (TIDE-sequencing). Briefly, genomic DNA (gDNA) was extracted from cells nucleofected with sgRNAs targeting *ROSA26, RFK* sgRNA #1 or sgRNA #2 at 96 hours post nucleofection. 250 ng of gDNA was combined with 10 μM primer sequences which amplify the genomic region targeted by sgRNAs (see Table S7 for sequence information), and the NEBNext High-Fidelity 2x PCR Master Mix. For amplifying the region targeted by sgRNA #1, the following PCR thermocycler conditions were used: Initial denaturation, 95ºC for 5 minutes; [Denaturation, 95ºC for 30 seconds; Annealing, 55ºC for 30 seconds; Extension, 72ºC for 1 minute], repeated for 34 cycles; Final extension, 72ºC for 5 minutes; Infinite hold, 4ºC. For amplifying the region targeted by sgRNA #2, the following PCR thermocycler conditions were used: Initial denaturation, 95ºC for 5 minutes; [Denaturation, 95ºC for 30 seconds; Annealing, 52ºC for 30 seconds; Extension, 72ºC for 1 minute], repeated for 34 cycles; Final extension, 72ºC for 5 minutes; Infinite hold, 4ºC. The PCR product was purified using the QiaQuick PCR purification kit and submitted for Sanger sequencing. The Synthego Inference of CRISPR Edits (ICE) platform was used to characterize edited alleles (https://ice.editco.bio/). Greater than 75% editing was achieved in all experiments.

### Nucleotide Supplementation Rescue Experiments

Cas9-competent MV4-11 cells harboring doxycycline-inducible sgRNAs against *ROSA26* or *RFK* #1 were induced with 1 μg/mL doxycycline for 96 hours. Cells were then seeded at a concentration of 0.2 × 10^6^ cells/well of a 12-well plate in 2 mL of Plasmax or RPMI medium in triplicate. 100 mM stock solutions of sterile-filtered purine (adenosine and guanosine) and pyrimidine (cytidine and thymidine) cocktails in H_2_O were prepared and pH adjusted to ensure complete solubility. Nucleoside stocks were supplemented into culture such that the final concentration of purine cocktail was 200 μM and pyrimidine cocktail was 500 μM. Supplementation experiments using cytidine or 2’-deoxycytidine hydrochloride alone were also performed by preparing sterile 100 mM stock solutions in H_2_O and supplementing into culture for a final concentration of 500 μM. After an additional 96 hours of culture in the presence of supplemented nucleotides, cell numbers were determined via CountBright Absolute Counting Beads and flow cytometry as described above. For validation of the CRISPR screen, the above conditions were repeated using NB4 cells in Plasmax complete or riboflavin deficient medium.

### Aconitase Assay

The aconitase assay was performed with the Abcam Aconitase Activity Assay Kit. Cas9-competent NB4 cells already transduced and sorted for 100% positive doxycycline-inducible sgRNAs tagged with GFP targeting *ROSA26* or *RFK*#1 were induced with 1 μg/mL doxycycline for 96 hours and expanded in triplicate. At day 9, 3 × 10^6^ cells were harvested for each condition, and resuspended in 100 μL Aconitase Preservation Solution with detergent. Cells were vortexed and incubated for 30 minutes on ice and then centrifuged at 20,000 *g* for 10 minutes at 4ºC. We performed a bicinchoninic acid (BCA) assay to quantify protein supernatant from each sample. 100 μg of each sample was then made up to 50 μL in Buffer and loaded to a well of the 96-well assay plate. 200 μL of Assay Buffer (including diluted isocitrate and manganese) was then added. A Buffer/Assay Buffer only background control was included. A ClarioStar PLUS plate reader was used for room-temperature colorimetric detection at OD 240 nm for 45 minutes, at an interval of 45 seconds with 3 second auto-shake between readings. Data are reported as mOD per minute per μg of protein.

### May-Grünwald Giemsa Staining

0.1 × 10^6^ cells in 100 μL of PBS per condition were spun onto a frosted glass slide at 300 *g* for 5 minutes using a Shandon Cytospin 4. After drying for 30 minutes at room temperature, slides were submerged in pure methanol for 5 minutes to allow for fixing and permeabilization and allowed to dry at room temperature. Slides were then submerged in May-Grünwald Stain for 5 minutes and washed in PBS for 5 minutes. Slides were then stained in Giemsa solution (1:20 diluted in H_2_O) for 20 minutes. Slides were rinsed with distilled water and dried at room temperature, prior to mounting with Shandon slide mounting resin and cover slipping. Slide images were captured via light microscopy.

### Synergy Experiments

For *RFK*-knockout synergy with venetoclax, Cas9-competent cells already transduced and sorted for 100% positive doxycycline-inducible sgRNAs tagged with GFP targeting *ROSA26* or *RFK*#1 were induced with 1 μg/mL doxycycline for 96 hours. 7,500 cells/well of a 384-well white-bottom plate in 50 μL of medium were seeded and treated with DMSO or escalating concentrations of venetoclax with three replicates for each concentration using a HP D300e Digital Dispenser. After 72 hours, 10 μL of CTG was added to each well with gentle agitation for 15 minutes prior to detection of luminescence with a ClarioStar PLUS plate reader. For analysis, background well readings were subtracted from each well, and cell viability expressed as a ratio of signal for each venetoclax-treated well relative to DMSO. For exogenous depletion of riboflavin and synergy with venetoclax, cells were cultured in Plasmax complete or Plasmax minus riboflavin for 4 days prior to the assay, which was otherwise performed as above.

For synergy experiments between roseoflavin and venetoclax, parental cells were seeded and treated with either DMSO or escalating concentrations of both roseoflavin and venetoclax as above. Viability data was captured via CTG as above. Synergy was determined by utilizing the SynergyFinder R package (v3.0) using the Bliss synergy model, for which a Bliss Synergy Index was computed for each cell line.

### Western Immunoblotting

Cells treated/induced as outlined were lysed in Laemmli lysis buffer (for 2X: 4 mL 10% SDS (4%); 2 mL Glycerol (20%); 1.2 mL 1M Tris-HCl pH 6.8; 2.8 mL ddH_2_O), vortexed vigorously, and boiled at 95ºC for 10 minutes. Lysates were quantified using a BCA assay, normalized, and sample buffer (containing 0.1% Bromophenol Blue in 2-mercaptoethanol) added 1:10 to each sample. Lysates were boiled for 10 minutes and then separated on 4-12% graded SDS-PAGE gels and transferred to a methanol-activated polyvinylidene difluoride (PVDF) membrane by electroblotting.

Membranes were then blocked in 5% skim milk diluted in Tris-buffered saline with 0.1% Tween-20 detergent (TBS-T) for 1 hour and then incubated in primary antibody (diluted in TBS-T + 5% bovine serum albumin (BSA) overnight at 4ºC with mild agitation. Membranes were then washed in TBS-T and incubated in appropriate secondary antibody (diluted in 5% skim milk) for 1 hour at room temperature. After washing, blots were developed on X-ray film using enhanced chemiluminescence prime (ECL Prime) development solution. For multiple probing on the same membrane, membranes were incubated for 30 minutes in Restore Western Blot Stripping Buffer, re-blocked with 5% skim milk for 1 hour at room temperature and then re-probed with alternate primary antibody overnight as described above.

### Mitochondrial Isolation and Blue Native (BN)-PAGE

Cas9-competent NB4 or MV4-11 cells harboring doxycycline-inducible sgRNAs against *ROSA26* or *RFK* #1 were induced with 1 μg/mL doxycycline for 96 hours and expanded. At day 9, 3 × 10^7^ cells per condition were harvested, washed twice in ice-cold PBS and snap frozen. Similarly, NB4 and MV4-11 cells were cultured for 4 or 7 days in Plasmax complete or riboflavin deprived culture medium, and media was refreshed every 48 hours until endpoint. At day 4 and day 7, 3 × 10^7^ cells per condition were harvested, washed twice in ice-cold PBS and snap frozen. Mitochondria were isolated from cells using a differential centrifugation protocol. Cells were resuspended in 4 mL of mitochondrial isolation buffer (20 mM HEPES-KOH, pH 7.6, 200 mM mannitol, 70 mM sucrose, 1 mM EDTA, 1 × protease inhibitor cocktail, 2 mg/mL BSA), homogenized via five strokes with a 15 mL dounce homogenizer, and the resultant lysate was centrifuged at 4 °C and 1000 *g* for 10 minutes to remove nuclear debris and intact cells. The supernatant, which contains mitochondria, was split across 2 × 2 mL tubes and centrifuged at 4 °C and 12000 *g* for 10 minutes to obtain crude mitochondrial pellets. Crude mitochondrial pellets for each sample were resuspended in 200 μL mitochondrial isolation buffer (lacking BSA), collected in a single 1.5 mL tube, and re-pelleted through centrifugation at 4 °C and 12000 *g* for 5 minutes. The final crude mitochondrial pellets were resuspended in 150-200 μL of mitochondrial isolation buffer (lacking BSA), and protein concentration was determined for each sample using a BCA assay. 100 μg or 50 μg aliquots of crude mitochondria were pelleted through centrifugation at 4 °C and 12000 *g* for 5 minutes and snap frozen following removal of the supernatant. For BN-PAGE, 100 μg aliquots of crude mitochondria were resuspended in 40 μL of NativePAGE Sample Buffer containing 1 x protease inhibitor cocktail and 1.5 % (w/v) digitonin (6 g/g digitonin/protein ratio). Samples were incubated on ice for 20 minutes before centrifugation at 18,000 *g* and 4 °C for 20 minutes to pellet insoluble debris. Clarified supernatants were transferred to clean 1.5 mL tubes containing 3 μL NativePAGE 5% G-250 sample additive immediately prior to loading. Samples (9 μL loaded per sample) were separated on 4-16% NativePAGE Bis-Tris Mini protein gels and transferred onto methanol activated PVDF membranes by electroblotting. Western blotting was then performed as described in the western immunoblotting section, except that blots were developed using the Amersham Imager 680 and SuperSignal West Pico or Femto chemiluminescent substrates. For SDS-PAGE analysis of crude mitochondria, 50 μg pellets were resuspended in NuPAGE LDS Sample Buffer, heated at 70 °C for 10 minutes, and separated on 4-12% NuPAGE Bis-Tris mini protein gels. Western transfer and immunoblotting were then performed the same as for BN-PAGE.

### Tandem Mass Tag Proteomics

Cas9-competent NB4 cells harboring doxycycline-inducible sgRNAs against *ROSA26* or *RFK* #1 were induced with 1 μg/mL doxycycline for 96 hours and expanded, or concurrently, cells with sgRNAs targeting *ROSA26* transitioned in Plasmax complete/Plasmax without riboflavin medium. At day 9 for genetic deletion of *RFK*, or day 4 for exogenous riboflavin depletion, 1 × 10^7^ cells per condition were harvested, washed twice in ice-cold PBS, and snap frozen. Cells were lysed and protein quantified, and 150 μg of protein from each sample was reduced with tris(2-carboxyethyl)phosphine (TCEP), alkylated with iodoacetamide, and further reduced with fresh dithiothreitol (DTT). Proteins were then precipitated onto SP3 beads to facilitate buffer exchange into digestion buffer, and samples subsequently digested with Lys-C (1:50 diluted) overnight at room temperature and trypsin (1:50 diluted) for 6 hours at 37ºC. Peptides were then labelled with TMTPro reagents, and 2 μL of each sample was pooled and used to shoot a ratio check to confirm complete TMT labelling. All TMTPro-labelled samples from each group were pooled and desalted by Sep-pak. Whole proteome peptides were fractionated into 24 fractions using basic reverse-phase high performance liquid chromatography (HPLC). 12 fractions were solubilized, desalted by stage tip, and analyzed on an Orbitrap Lumos mass spectrometer.

The output MS2 spectra were searched using the COMET algorithm against a Human Uniprot composite database containing its reversed complement and known contaminants. Peptide spectral matches were filtered to a 1% false discovery rate (FDR) using the target-decoy strategy combined with linear discriminant analysis. The proteins were filtered to a <1% FDR and quantified only from peptides with a summed SN threshold of >160. The search parameters are outlined below:

16-plex Comet Search Parameters:

- Peptide Mass Tolerance: 50 ppm
- Fragment Ion Tolerance: 0.4 ppm
- Max Internal Cleavage Site: 2
- Max differential/Sites: 5
- Methionine oxidation is used as variable modification.

16-plex TMTpro Reporter Quant Parameter:

- Proteome (MS3) - tolerance = 0.003, ms2_isolation_width = 0.7, ms3_isolation_width = 1.2 Summed intensity and normalized relative abundance values (%) were calculated for each protein and in each condition and subsequently used to determine log_2_ fold changes in abundance and significance between conditions.

### LC/MS-MS Metabolite Profiling

For untargeted metabolite profiling, Cas9-competent NB4 cells harboring doxycycline-inducible sgRNAs against *ROSA26* or *RFK* #1 were induced with 1 μg/mL doxycycline for 96 hours in RPMI medium and expanded in biological quadruplicate with media changes every 48 hours. At day 9, 10 × 10^6^ cells per condition were harvested, washed in ice-cold blood bank saline, and metabolites extracted in 1 mL ice-cold extraction buffer (40:40:20 acetonitrile:methanol:water, containing ^13^C/^15^N-labelled amino acid standards and ^13^C-labelled riboflavin standard). Cell suspension was vortexed vigorously for 10 minutes at 4ºC and then centrifuged at 20,000 *g* for 10 minutes at 4ºC. Metabolite extract was removed and dried under nitrogen gas and resuspended in 100 μL LC/MS-grade water. Untargeted metabolite profiling was conducted on a QExactive HF-X mass spectrometer. The analytes were ionized via a HESI II probe. The mass spectrometer was coupled to a Vanquish binary UPLC system (Thermo Fisher Scientific). Two chromatographic separations were performed for each sample; in both, 5 μL of sample was injected into a BEH Z-HILIC column (100 × 2.1 mm, 1.7 μm). First separation was carried out with 15 mM ammonium bicarbonate in 90% water and 10% acetonitrile as mobile phase A, and mobile phase B was 15mM ammonium bicarbonate in 95% acetonitrile and 5% water. The chromatographic gradient was carried out at a flow rate of 0.225 ml/min and was adapted from Koley, et al^87^. The mass analysis was carried out in negative mode. The second separation was adapted from Mülleder et al^88^. Both mobile phases A and B were buffered with 10 mM ammonium formate and 45 mM formic acid (pH 2.7). In mobile phase A, the solvents were in 1:1 acetonitrile:water. Mobile phase B consisted of buffer in 95:5:5 acetonitrile:water:methanol. The column oven was held at 40°C and autosampler at 4°C. The chromatographic gradient was run at a flow rate of 0.4 ml/min as follows: 0.75 min initial hold at 95% B; 0.75-3.00 min linear gradient from 95% to 30% B, 1.00 min isocratic hold at 30% B. B was brought back to 95% over 0.50 minutes, after which the column was re-equilibrated under initial conditions. Metabolites were analyzed in positive mode. Sample acquisition was conducted in full-scan mode, with the spray voltage set to 3 kV (negative) or 3.5 kV (positive), the capillary temperature to 320°C, and the HESI probe to 300°C, sheath gas to 40U, the AUX gas to 8U, and the sweep gas to 1 unit, and resolving power to 120,000. An untargeted metabolite library was created via top-15 DDA acquisitions on a pooled study sample. MS1 resolution was set to 60,000 and MS2 to 30,000. The .raw data files were processed in MZmine^89^ and emzed^90^. Metabolite MS2 spectra were compared to HMDB, GNPS, Massbank, and MoNA databases using “Spectral library search”. In addition, retention times and m/z values of a full-scan acquisition on a pooled study sample was compared to an in-house database of retention times determined for authentic standards (Human Endogenous Metabolite Compound Library Plus). Internal standards were integrated using emzed, and raw peak areas were normalized by dividing by sample biomass and mean internal standard area.

For flavin measurements in AML cells, samples were prepared as above; for flavin measurement in blood plasma, harvested whole blood was collected using EDTA vacutainers and then centrifuged at 2000 *g* at 4ºC for 20 minutes. Plasma was transferred to a new tube. 40 μL of blood plasma was used for each sample, and metabolites were extracted in 1 mL ice-cold extraction buffer (100% LC/MS-grade methanol with 13C-labelled riboflavin standard). Metabolite extract was dried under nitrogen gas and resuspended in 100 μL LC/MS-grade water. 5 μL of each sample was injected into a Kinetex XB-C18 column (particle size 1.7 μm, 50 × 2.1 mm; 100 Å); a gradient was applied as described previously^91^. Samples were run on a Vanquish binary UPLC system coupled with a Q-Exactive HF-X mass spectrometer. The mass spectrometer was operated in full scan positive mode at mass resolution of 120,000 for quantification; for identification, PRM was applied at MS2 resolution 30,000 and N(CE) 35, 50, and 150. Heated electro-spray ionization probe was used applying the following source parameters: vaporizer 300°C; ion spray voltage +3.50 kV, sheath gas, 50; aux gas 10; sweep gas, 1; S Lens RF level, 35.0; capillary temperature, 320°C, scan range m/z 300-900. Peak area for each analyte was determined via linear integration using emzed^90^ and normalized to the peak area of labelled riboflavin standard.

For nucleotide metabolite profiling, Cas9-competent NB4 cells harboring doxycycline-inducible sgRNAs against *ROSA26* or *RFK* #1 were induced with 1 μg/mL doxycycline for 96 hours in RPMI medium and expanded in biological quadruplicate with media changes every 48 hours. At day 7, 3 × 10^6^ cells per condition were harvested, washed in ice-cold blood bank saline, and metabolites extracted in 1 mL ice-cold extraction buffer (40:40:20 acetonitrile:methanol:water, 0.1M formic acid, containing ^13^C/^15^N-labelled amino acid standards). Cell suspension was vortexed vigorously for 10 minutes at 4ºC and then centrifuged at 20,000 *g* for 10 minutes at 4ºC. Metabolite extract was removed and dried under nitrogen gas and resuspended in 100 μL LC/MS-grade water. Metabolites were measured using a Dionex UltiMate 3000 ultra-high performance liquid chromatography system connected to a QExactive benchtop Orbitrap mass spectrometer, equipped with an Ion Max source and a HESI II probe (Thermo Fisher Scientific). External mass calibration was performed using the standard calibration mixture every 7 days. Samples were separated by chromatography by injecting 2 μl of sample on a SeQuant ZIC-pHILIC Polymeric column (2.1 × 150 mm 5 μm). Flow rate was set to 150 μl min, temperatures were set to 25°C for column compartment and 4°C for autosampler sample tray. Mobile Phase A consisted of 20 mM ammonium carbonate, 0.1% ammonium hydroxide. Mobile Phase B was 100% acetonitrile. The mobile phase gradient was as follows: 0–20 minutes: linear gradient from 80% to 20% B; 20–20.5 minutes: linear gradient from 20% to 80% B; 20.5–28 minutes: hold at 80% B. Mobile phase was introduced into the ionization source set to the following parameters: sheath gas = 40, auxiliary gas = 15, sweep gas = 1, spray voltage = 3.1 kV, capillary temperature = 275°C, S-lens RF level = 40, probe temperature = 350°C. Metabolites were monitored in full-scan, polarity-switching, mode. An additional narrow range full-scan (220–700 m/z) in negative mode only was included to enhance nucleotide detection. The resolution was set at 70,000, the AGC target at 1 × 10^6^ and the maximum injection time at 20 msec. Relative quantitation of metabolites was performed with XCalibur using a 5-ppm mass tolerance and referencing an in-house retention time library of chemical standards. Raw peak areas of metabolites were median normalized by sample after an initial normalization to the abundance of internal ^13^C/^15^N-labelled amino acid standards.

### Seahorse Oxygen Consumption Experiments

Cas9-competent NB4 and MV4-11 cells harboring doxycy-cline-inducible sgRNAs against *ROSA26* or *RFK* #1 were induced with 1 μg/mL doxycycline for 96 hours and expanded. Agilent XFe96 cartridges were hydrated in calibration solution overnight the day before the experiment as per the manufacturer’s instructions, and Agilent XFe96 cell culture plates were coated the day before with Cell-Tak to allow for suspension cell adhesion. At day 9, 0.1 × 10^6^ cells were seeded in XFe96 cell culture plates using RPMI culture medium with 1 mM sodium pyruvate at pH 7.2. The assay measured oxygen consumption rate at basal conditions and then upon sequential injections of oligomycin (1.5 μM), carbonyl cyanide p-(trifluoromethoxy) phenylhydrazone (FCCP) (2 μM), and rotenone/antimycin (2 μM). After each injection, three measurements were obtained. At least 6 replicates were utilized to calculate each data point.

### RNA-Sequencing and Analysis

Cas9-competent NB4 cells harboring doxycycline-inducible sgRNAs against *ROSA26* or *RFK* #1 were induced with 1 μg/mL doxycycline for 96 hours and expanded, with media refreshed every 48 hours. At day 7, 3 × 10^6^ cells were harvested, washed in PBS, and total RNA extracted from cells with on-column DNA digestion. Sequencing libraries (poly-A capture, non-directional library preparation) were generated by Novogene according to the manufacturer’s protocol, and were sequenced on the Illumina NovaSeq 6000, with paired end 150 bp reads (Q30 ≥ 85%) to a depth of 20 million reads per sample. For each experiment, technical triplicates were used for each condition. Reads were mapped to the GRCh38/hg38 human genome using STAR v2.7^81^. Quality control tests for mapped reads and for replicate reproducibility were performed using SAR-Tools (v1.7.4)^83^. Gene-level counting of reads was conducted using FeatureCounts (v1.6.3), a component of the subread v2.0.0 package^82^. Gene counts were normalized and used to quantify differentially expressed genes between the experimental and control conditions using the DESeq2 (v1.34.0) method implemented in Bioconductor v3.9^84^. Genes with ≥ 10 reads across at least three samples were annotated as expressed. Differentiability for expressed genes was assessed with DESeq2 based on the robust shrunken log_2_ fold change scores and the approximate posterior estimation for GLM coefficients (apeglm v1.16.0^85^; method for effect size). All RNA-seq analysis figures were generated using R (v4.1.1) software, and the cut-offs for DEGs is described in each figure. GO Term analysis was performed using ToppGene software using term sets as described^86^.

### Generation of Metabolic CRISPR-Cas9 Library

Metabolism associated genes from previously published CRISPR-Cas9 screening libraries^19,48,92^ and the Mammalian Metabolic Enzyme Database^93^ were collated, converted into human gene equivalents (in the context of mouse genes as appropriate), and replicates removed. We filtered the gene list and explicitly removed non-metabolic genes, cell cycle-related genes, epigenetic enzymes, receptors, transcription factors and ribosomal genes. Two Cas9-optimized sgRNAs targeting each gene were designed, using the Broad Institute’s CRISPick sgRNA design platform (publicly available at https://portals.broadinstitute.org/gppx/crispick/public). We additionally included 38 positive control genes; 28 genes whose deletion has been shown to induce AML cell differentiation and 10 common essential genes. 250 negative control non-targeting sgRNAs targeting gene deserts were also included. sgRNA sequences and genes targeted by the library are included in **Table S4**. Upon synthesis of sgRNAs, the library was assembled by cloning sgRNAs into the pXPR-050 vector that expresses a puromycin resistance cassette as previously described^94^.

### CRISPR-Cas9 Screens

For the screen performed in riboflavin free culture medium, Cas9-competent NB4 cells were transduced with the metabolic CRISPR knockout library in two biological replicates at an MOI of ~0.3-0.4. 24 hours post infection, cells were selected with 1 μg/mL puromycin for a further 72 hours. After confirming MOI was within the target range, cells were counted and split into two sets of >6.072 × 10^6^ cells per replicate to maintain library representation of 1000X or greater: one in Plasmax complete medium and the other in Plasmax medium with no riboflavin. Cells were counted and passaged every 4 days, and media was refreshed every 48 hours to prevent media exhaustion. After 14 days of culture, cells were harvested and washed in PBS. Genomic DNA was extracted from the cell pellet using the NucleoSpin Blood L kit. sgRNA sequences were then PCR amplified and submitted for standard Illumina sequencing^95^.

For screen analysis, the sgRNAs read count data for both cell lines were deconvoluted from sequence reads by using the Broad Institute’s PoolQ software (https://portals.broadinstitute.org/gpp/public/software/poolq). The sgRNA reads were then log-normalized to the initial plasmid pool (pDNA). Quality control pre-processing steps were performed to remove guides with low efficacy (<0.50) and the guides with low representation in the initial plasmid pool (pDNA). The sgRNA guides were then mapped to genes based on the previously defined annotations, and positive/negative controls (defined above) were used in further downstream analysis. The sgRNA dependency data log-normalized to pDNA were processed on the Genomic Perturbation Platform at the Broad Institute. Gene level dependency scores were computed from the sgRNA log-norm data by using the hypergeometric distribution tool (https://portals.broadinstitute.org/gpp/public/). The magnitude of the gene-level dependency effect was inferred as the average log-norm dependencies of the sgRNAs assigned to the gene. The *P*-values were computed using the probability mass function of a hypergeometric distribution for the log-norm dependency-ranked sgRNAs. Significance was estimated based on the cut off absolute (average log-fold change) > 0.5 and average (−log10(*P*-value)) > 1.

### *In Vivo* Studies

For riboflavin starvation experiments in non-tumor bearing animals, 10 NSG mice were randomly assigned to either a riboflavin competent (0.006 g/kg) or deficient diet for 3 weeks, with five mice per group. Mouse mass measurements were taken every 4 days, and no mice reached ethical end-point from riboflavin starvation, which included pain and distress (hunched and ruffled coats); labored movement; and being moribund. At planned takedown, all mice were sacrificed using CO_2_ asphyxiation, and whole blood immediately captured via cardiac puncture using EDTA vacutainers. Blood was centrifuged at 2000 *g* for 20 minutes at 4ºC, and plasma supernatant captured for downstream metabolite profiling. One mouse was censored in the riboflavin-deficient arm of the study due to dehydration caused by a faulty water dispenser.

For the cross-sectional study testing anti-tumor effects of *RFK* KO in AML, 0.25 × 10^6^ Cas9-competent MV4-11 cells harboring doxycycline-inducible sgRNAs against *ROSA26* or *RFK* #1 were injected into randomly assigned NSG mice (*n* = 8 mice per group). Engraftment of leukemias was determined via bone marrow aspiration weekly until 1% human CD45-positive cells were detected via flow cytometry (14 days). Upon confirmation of engraftment, mice were placed on doxycycline containing chow (625 ppm) to induce sgRNAs and leukemia progression allowed to continue for 14 additional days after which mice were sacrificed using CO_2_ asphyxiation, and bone marrow, dissociated spleen and peripheral blood were collected. Leukemic burden was assessed in each compartment by the ratio of mouse CD45-APC to human CD45-V450 cells as determined by flow cytometry. One mouse was censored from the sgRosa group of the study due to complications from bone marrow aspiration.

### DepMap Analyses

Normalized dependency scores (Chronos), and mRNA expression log_2_(TPM+1) data for *RFK* were downloaded from the DepMap Portal data downloads section for all cell lines screened in release 24Q2 (https://depmap.org/portal/data_page/?tab=customDownloads). Dependency and expression plots were presented as Chronos scores across lineage and, within each lineage, specific disease subtypes. For genetic co-dependency analysis with *RFK*, a custom analysis was performed using Pearson correlation between CRISPR dependency of *RFK* and gene expression across all other genes in every cell line (*n* = 1,150). The same Pearson correlation was performed using RFK dependency and metabolomics data.

### Statistics

Data for each figure are presented as mean ± SD unless otherwise stated in the Figure Legends. Statistical tests are detailed in the corresponding figure legends and are calculated using GraphPad Prism v10.4, R v4.4.1 or Microsoft Excel v16.80. Figures were generated using either GraphPad Prism or R. No data were excluded.

---

**Figure S1.**
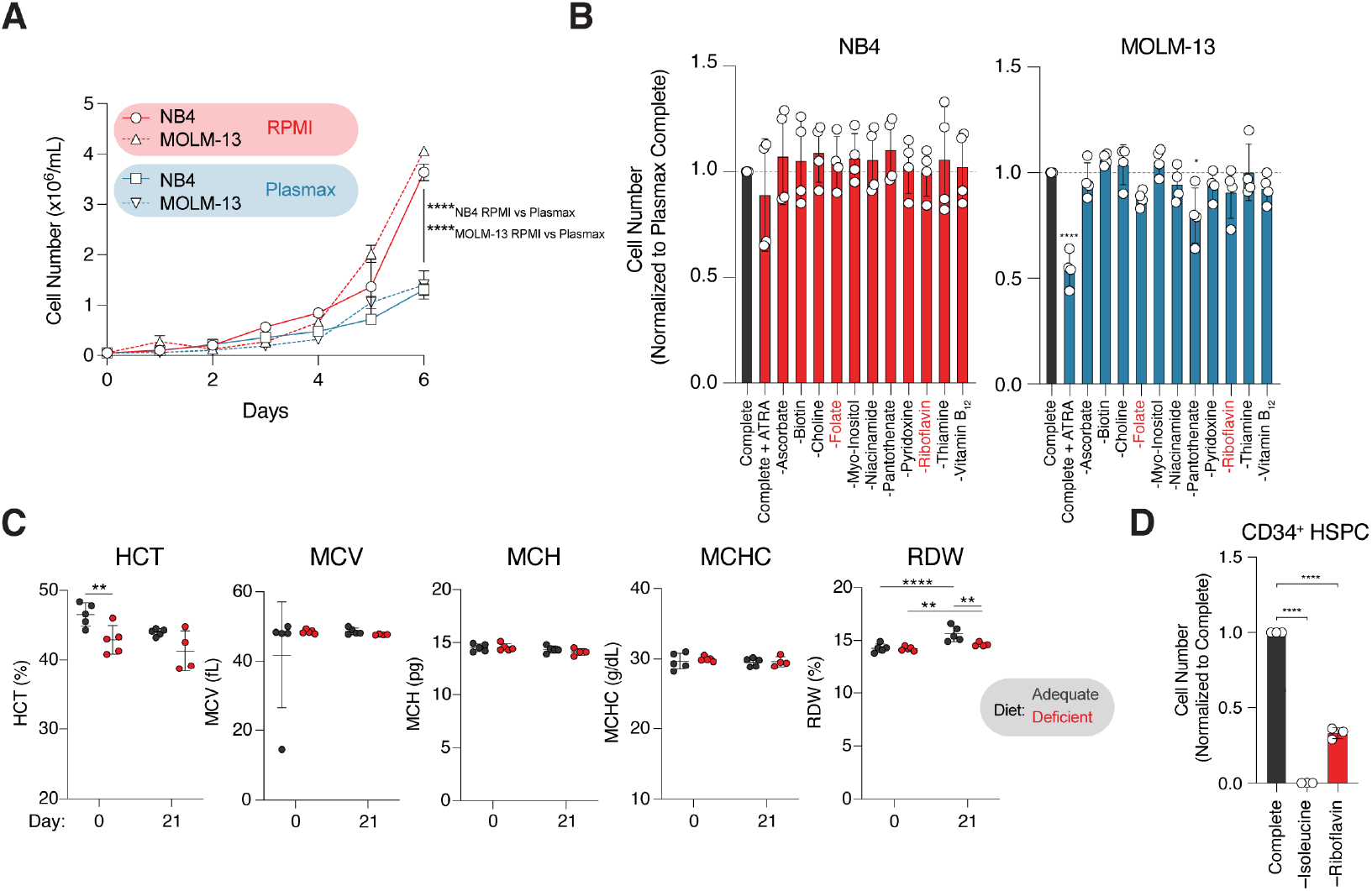
Physiologically relevant metabolite depletion screens identify riboflavin as a leukemic dependency. **A**. Proliferation of NB4 and MOLM-13 cells in standard RPMI culture conditions and Plasmax complete over 6 days. No media changes were performed. Data representative of *n*=3 biological replicates. **B**. Proliferation of NB4 and MOLM-13 cells upon systematic removal of vitamins from Plasmax medium at day 3. Cell number is normalized to Plasmax complete medium. Cells cultured in Plasmax complete medium and treated with 1 μM all-*trans* retinoic acid (ATRA) served as an anti-proliferation/differentiation control. Data representative of *n*=4 pooled replicates from two independent biological experiments. **C**. Complete blood counts of mice fed adequate versus deficient riboflavin diets at start of experiment (baseline, day 0) and at 21 days. HCT, hematocrit test; MCV, mean corpuscular volume test; MCH, mean corpuscular hemoglobin test; MCHC, mean corpuscular hemoglobin concentration test; RDW, red cell distribution width test. **D**. Cell number of healthy CD34^+^ HSPC cells after 14 days of culture in Plasmax complete or Plasmax lacking the essential amino acid isoleucine (anti-proliferative control) or riboflavin. Cell number is normalized to Plasmax complete medium. Data representative of *n*=3 biological replicates. The experiment was repeated with two separate donors. Data are presented as the mean ± SD. \*\**P* < 0.01 and \*\*\*\**P* < 0.0001 by unpaired two-tailed Student’s t-test (A, C) and ordinary one-way ANOVA with Bonferroni’s multiple comparisons test (B, D).

**Figure S2.**
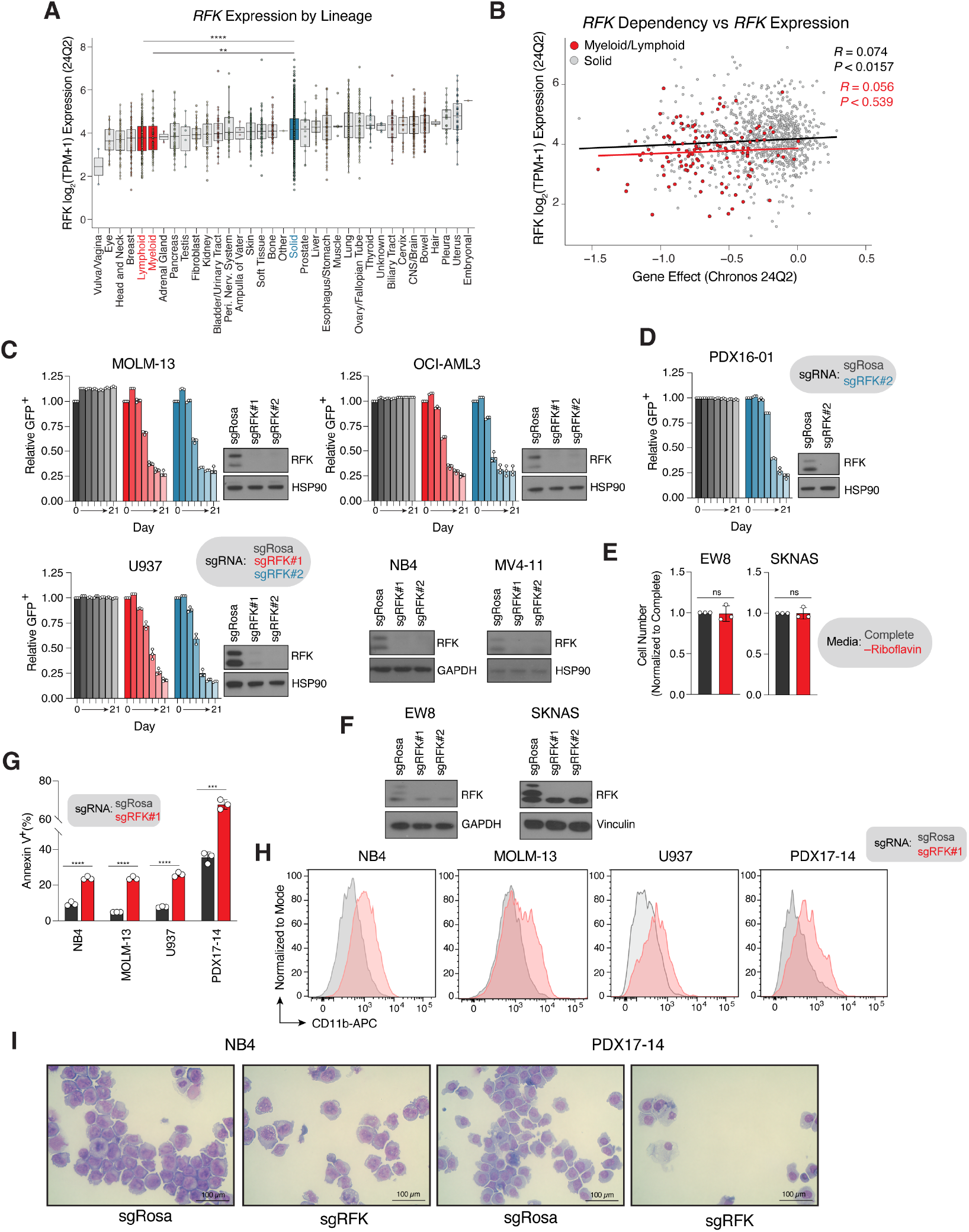
Riboflavin kinase (RFK) is a metabolic dependency in acute myeloid leukemia. **A**. Batch-corrected mRNA-level expression of *RFK* in cancer cell lines by disease lineage from the Broad Institute’s DepMap 24Q2 database. Expression reported as log_2_(TPM+1). Myeloid and lymphoid lineages are highlighted in red, with pooled solid cancer cell lines in blue. Boxplots ranked by median *RFK* expression. **B**. Correlation of *RFK* mRNA expression log_2_(TPM+1) versus *RFK* dependency (Gene Effect Chronos score). Each data point represents an individual cell line. Myeloid/lymphoid cell lines are highlighted in red, solid cancer cell lines in grey. Pearson correlation coefficients and significance values are reported for each. **C**. Competitive proliferation experiments in MOLM-13, OCI-AML3 and U937 AML cell lines over 21 days in RPMI medium. Two independent *RFK* sgRNAs and a control targeting the conserved *ROSA26* locus (sgRosa) were used, and relative proliferation was measured using GFP as a readout of edited cells. Data representative of *n*=3 biological replicates. Western immunoblot of RFK in NB4, MOLM-13, U937, OCI-AML3 and MV4-11 cells infected with a control sgRNA targeting Rosa or two independent sgRNAs targeting *RFK*. GAPDH (NB4) or HSP90 (all others) served as the loading control. **D**. Competitive proliferation experiments in Cas9-competent PDX16-01 cells subjected to short-term *in vitro* culture over 21 days in PDX medium. An *RFK* sgRNA and a control targeting Rosa was used, and relative proliferation was measured using GFP as a readout of edited cells. Data representative of *n*=3 biological replicates. Validation Western immunoblot of *RFK* KO shown, with HSP90 serving as the loading control. **E**. Cell number of EW8 Ewing sarcoma and SKNAS neuroblastoma cells upon exogenous riboflavin depletion using Plasmax medium at 14 days. Cell number is normalized to Complete medium. Data representative of *n*=3 biological replicates. **F**. Western immunoblot of RFK in EW8 and SKNAS cells infected with a control sgRNA targeting Rosa or two independent sgRNAs targeting *RFK*. GAPDH (EW8) or Vinculin (SKNAS) served as the loading control. **G**. Cell death in NB4, MOLM-13, U937 and PDX17-14 cells as measured by Annexin V-APC cell surface staining at day 9 (U937, PDX17-14) and day 11 (NB4, MOLM-13) post induction of Rosa or *RFK* sgRNAs. Data representative of *n*=3 biological replicates. **H**. Representative histograms of cell surface CD11b-APC in NB4, MOLM-13, U937 and PDX17-14 cells at day 9 (U937, PDX17-14) and day 11 (NB4, MOLM-13) post induction of Rosa or *RFK* sgRNAs. Data representative of two independent experiments. **I**. May-Grünwald Giemsa staining of NB4 and PDX17-14 cells after 9 days of induction of Rosa or RFK-targeting sgRNAs. Indicated scale on all images is 100 μm. Data are presented as the mean ± SD. \*\**P* < 0.01, \*\*\**P* < 0.001 and \*\*\*\**P* < 0.0001 by ordinary one-way ANOVA with Bonferroni’s multiple comparisons test (A), and individual unpaired two-tailed Student’s *t*-test (E, G).

**Figure S3.**
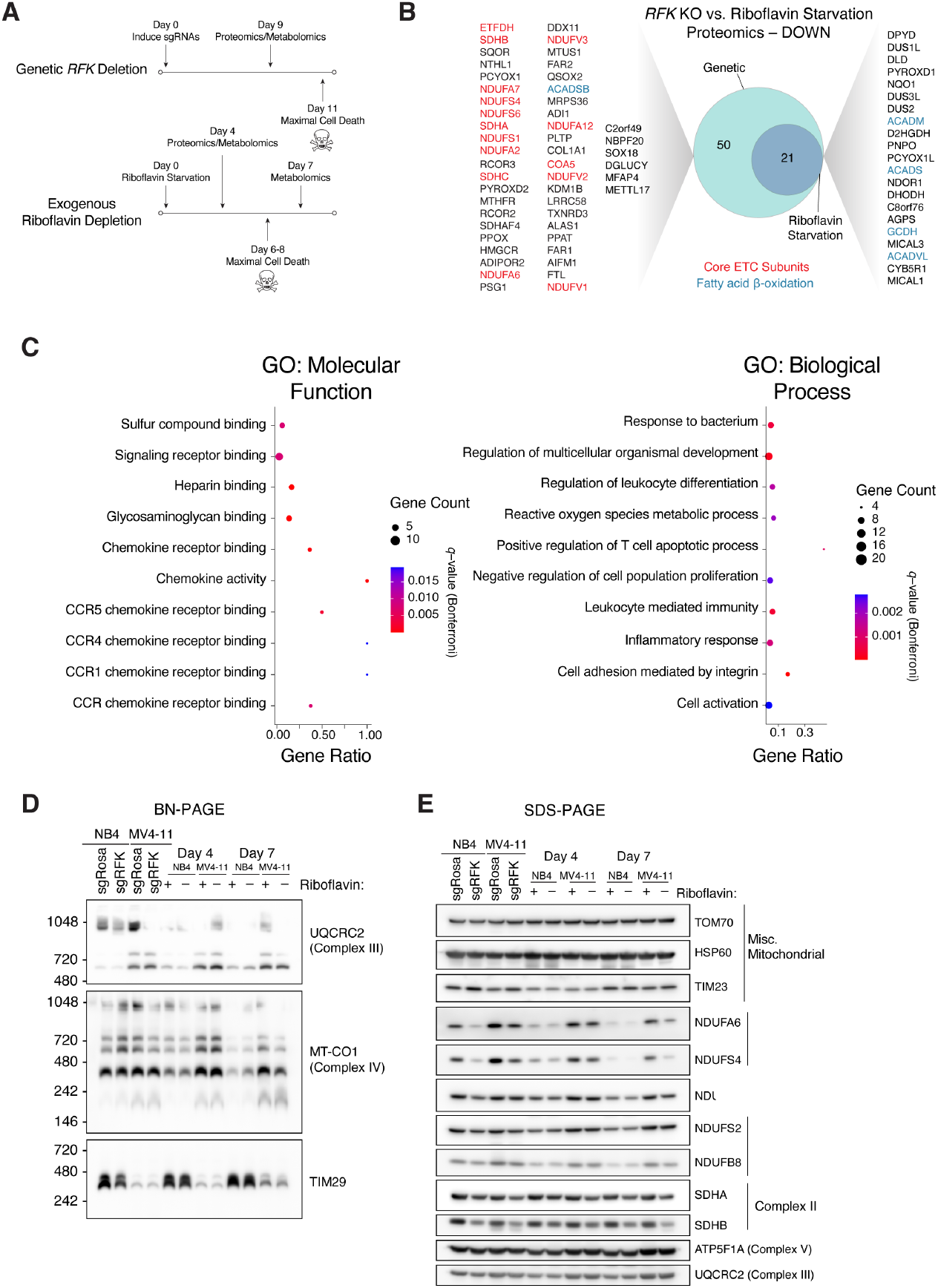
Global analysis of total proteome under riboflavin metabolism perturbation. **A**. Schema of proteomic/metabolomic experimental strategy. **B**. Venn diagram of differentially downregulated proteins upon exogenous riboflavin depletion (day 4, small circle) and genetic depletion of *RFK* (day 9, large circle). No unique proteins were detected in the exogenous riboflavin depletion condition. Proteins highlighted in red denote core electron transport chain subunits/components; blue denotes proteins associated with fatty acid β-oxidation. **C**. Bubble plots of gene ontology molecular function (left) and biological process (right) associated with differentially upregulated proteins from Figure 3B. *q*-values with Bonferroni correction are reported. **D**. Blue Native (BN) polyacrylamide gel electrophoresis (PAGE) of NB4 and MV4-11 cells at 9 days post deletion of *RFK* and after 4 and 7 days of exogenous riboflavin starvation. **E**. Sodium dodecyl sulfate (SDS) polyacrylamide gel electrophoresis (PAGE) of NB4 and MV4-11 cells at 9 days post deletion of *RFK*, and after 4 and 7 days of exogenous riboflavin starvation.

**Figure S4.**
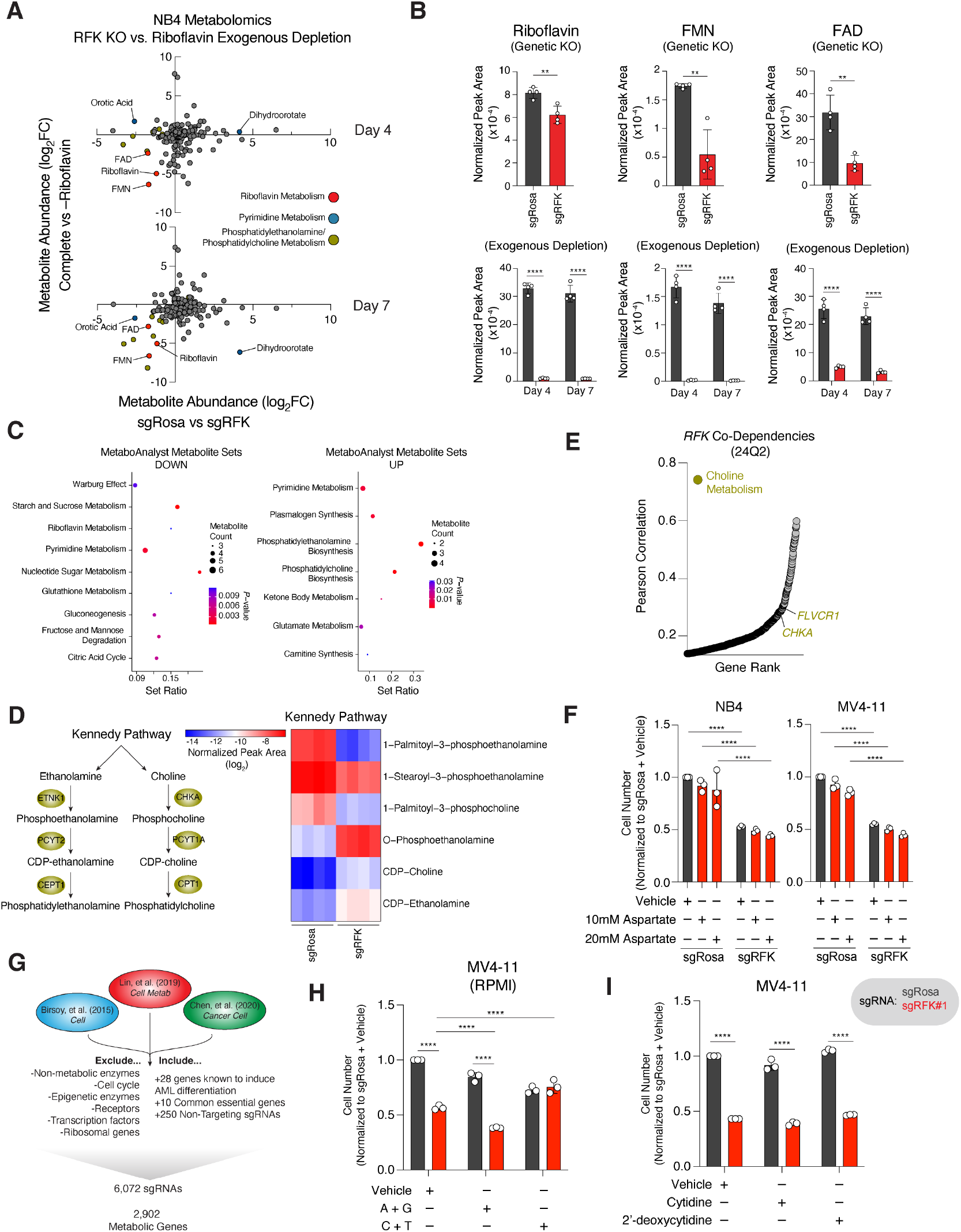
Riboflavin metabolism perturbation induces nucleotide imbalance. **A**. Dot plot of differentially altered metabolites upon exogenous depletion of riboflavin compared to complete medium (y-axis) at day 4 (top graph) and day 7 (bottom graph), versus deletion of *RFK* at day 9 (both graphs, *x-*axis). Relative log_2_ of metabolite abundances shown. Highlighted metabolites belong to the metabolic processes indicated. **B**. Quantification of riboflavin, FMN and FAD in NB4 cells upon 9 days of *RFK* knockout versus Rosa control via metabolite profiling (top) and in NB4 cells upon 4 and 7 days of exogenous riboflavin depletion versus complete medium (bottom). Normalized peak area of *n*=4 biological replicates for each condition shown. **C**. Bubble plots of gene ontology metabolite sets showing pathways associated with metabolites with decreased abundance (left) and increased abundance (right). *P*-values are reported. Pathway enrichment performed using MetaboAnalyst 6.0. **D**. Schema of the Kennedy Pathway (left) and heatmap of key pathway intermediates in NB4 cells after 9 days of *RFK* depletion versus Rosa control. Scale depicts normalized peak area of each metabolite. **E**. Dot plot of genetic co-dependency of genes with *RFK* ranked by Pearson correlation. Data from DepMap 24Q2. Yellow shading denotes choline metabolism-associated genes. **F**. Cell number of NB4 and MV4-11 cells at day 8 post induction of Rosa or *RFK* sgRNAs, treated with vehicle (water), or 10 mM or 20 mM aspartate in Plasmax medium for 4 days. Cell number normalized to sgRosa + Vehicle. Data representative of *n*=3 biological replicates. **G**. Schema of the design and generation of metabolism-focused CRISPR-Cas9 library. **H**. Cell number of MV4-11 cells at day 8 post induction of Rosa or RFK sgRNAs, treated with vehicle (water), or cocktails of purine (A+G, adenosine + guanosine) or pyrimidine (C+T, cytidine + thymidine) nucleosides in RPMI medium for 4 days. Cell number normalized to sgRosa + Vehicle. Data representative of *n*=3 biological replicates. **I**. Cell number of MV4-11 cells at day 8 post induction of Rosa or RFK sgRNAs, treated with vehicle (water), cytidine or 2’-deoxycytidine nucleosides in Plasmax medium for 4 days. Cell number normalized to sgRosa + Vehicle. Data representative of *n*=3 biological replicates. Data are presented as the mean ± SD. \*\**P* < 0.01, \*\*\*\**P* < 0.0001 by unpaired two-tailed Student’s *t*-test (B), and ordinary two-way ANOVA with Bonferroni’s multiple comparisons test (B, F, H, I).

**Figure S5.**
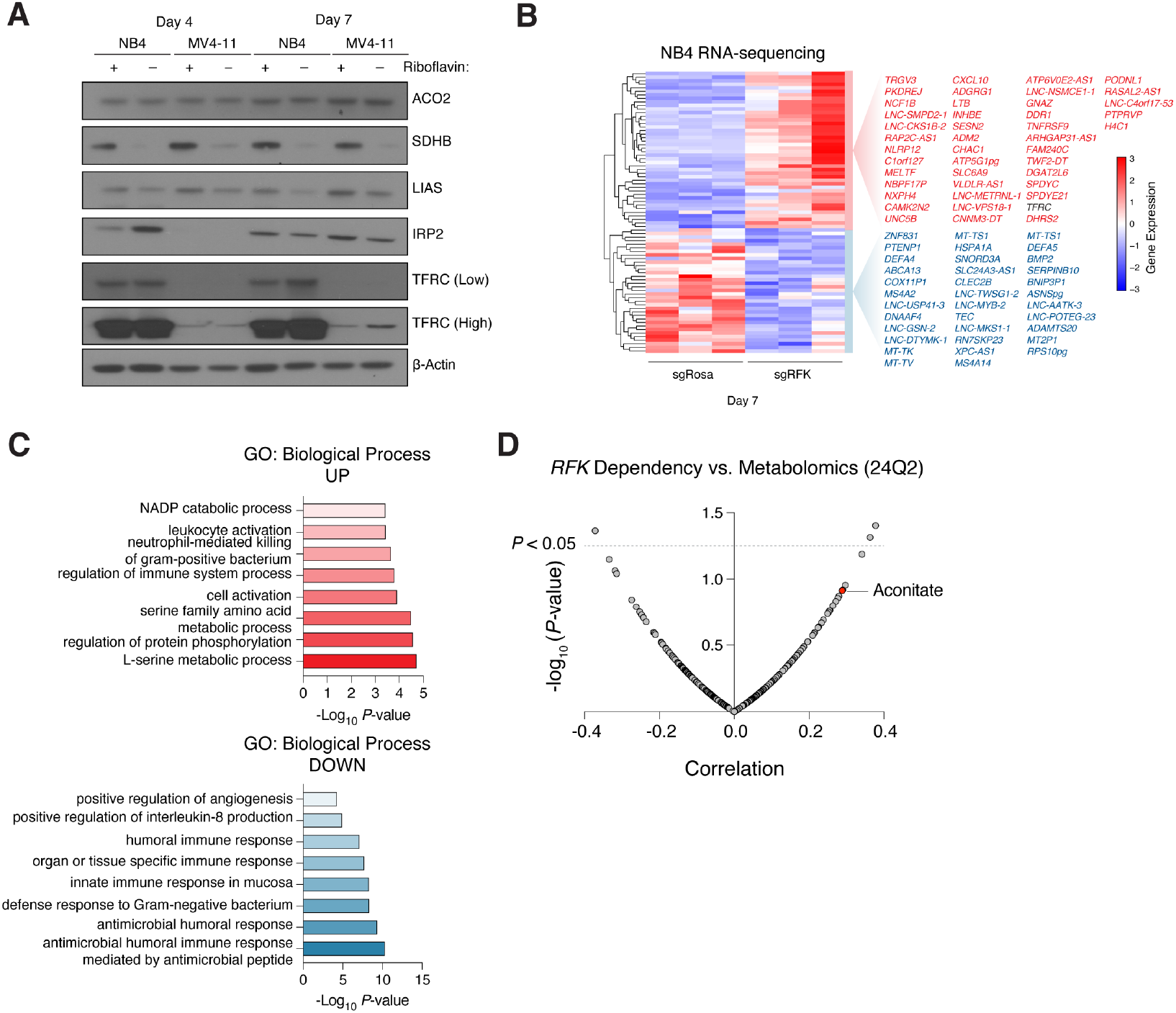
Riboflavin perturbation impairs iron-sulfur cluster dependent proteins. **A**. Western immunoblot analysis of indicated proteins in cell lysates isolated from NB4 and MV4-11 cells at days 4 and 7 post exogenous riboflavin withdrawal. β-Actin served as the loading control. Note the loading control is the same as Figure 4D, as this gel was stripped and re-blotted. **B**. Clustered heatmap of top differentially expressed genes in NB4 cells after 5 or 7 days of *RFK* depletion versus Rosa control. Upregulated genes are in red and downregulated genes are in blue. **C**. Bar graphs of enriched gene ontology biological process in upregulated genes (top) and downregulated genes (bottom) in NB4 cells with *RFK* deletion for 7 days. −log_10_(P-values) reported. **D**. Dot plot of the correlation between *RFK* dependency and metabolite abundance in the DepMap 24Q2 dataset ranked by significance. Each circle denotes an individual metabolite. The dotted line denotes *P* < 0.05.

**Figure S6.**
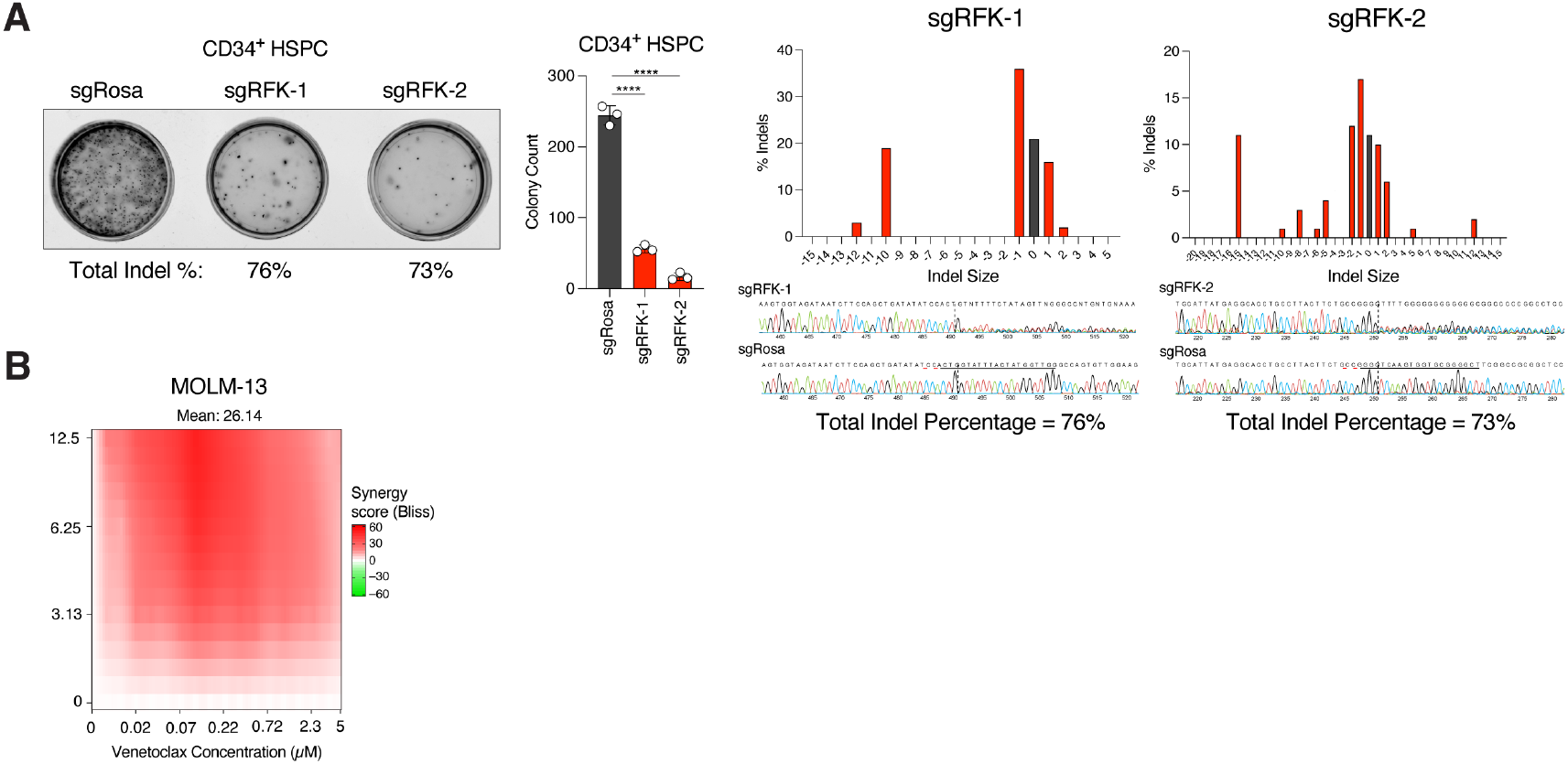
Perturbing riboflavin metabolism synergizes with BCL-2 inhibition. **A**. CD34^+^ HSPC cells nucleofected with recombinant Cas9 and sgRNAs targeting Rosa or with two independent sgRNAs targeting *RFK* for 15 days. Colony formation capacity was determined (left) and colony number was quantified (middle). Effective editing of *RFK* by the two sgRNAs was determined using Tracking of Indels by Decomposition (TIDE)-sequencing (right). **B**. MOLM-13 cells were treated with escalating concentrations of venetoclax and roseoflavin for 72 hours to determine viability effects. The presence of treatment synergy was determined using SynergyFinder and the Bliss synergy index and is denoted as regions of red in the graphs. The mean of three biological replicates was used for each data point. Data are presented as the mean ± SD. \*\*\*\**P* < 0.0001 by ordinary one-way ANOVA with Bonferroni’s multiple comparisons test.

